# Three Conserved Immune Dysfunction and Exclusion Subtypes in Bladder and Pan-cancers: Prognostic and Immunotherapeutic Significance

**DOI:** 10.1101/2023.10.18.562846

**Authors:** Kun Zheng, Youlong Hai, Hongqi Chen, Xiaoyong Hu, Kai Ni

**Author notes:** **Correspondence:** Kai Ni and Xiaoyong Hu Corresponding author Kai Ni will be responsible for handling correspondence at all stages of refereeing and publication. **Declarations of interest:** none. These authors contributed equally to this study (Kun Zheng, Youlong Hai, Hongqi Chen).

## Abstract

Molecular subtyping is expected to enable bladder cancer (BC) precise treatment. However, its clinical application remains defective and controversial. Given the significance of tumor immune dysfunction and exclusion (TIDE) in tumor immune escape and immunotherapy, we aimed to develop a novel TIDE-based subtyping method to facilitate personalized management. Transcriptome data of BC was used to evaluate the heterogeneity and the status of TIDE patterns. We identified 69 TIDE biomarker genes and classified BC samples into three subtypes. Subtype I showed the lowest TIDE status and malignancy with the best prognosis and highest sensitivity to immune checkpoint blockade (ICB) treatment, which was enriched of metabolic related signaling pathways. Subtype III represented the highest TIDE status and malignancy with the poorest prognosis and resistance to ICB treatment, resulting from its inhibitory immune microenvironment and T cell terminal exhaustion. Subtype II was in a transitional state with intermediate TIDE level, malignancy, and prognosis. We further confirmed the existence and characteristics of our novel TIDE subtypes using real-world BC samples. This subtyping method was proved to be more efficient than known methods in identifying non-responders to immunotherapy. We also found that combining our TIDE subtypes with known biomarkers can improve the sensitivity and specificity in predicting ICB response. Moreover, besides guiding ICB treatment, this classification approach can assist in selecting the frontline or recommended drugs. Finally, the TIDE subtypes are conserved across pan-tumors. In conclusion, our novel TIDE-based strategy is a powerful clinical tool for BC and pan-cancer patients, and potentially guiding personalized immunotherapy.

## Background

Bladder cancer (BC) is a prevalent malignancy of the urinary system, ranking ninth in incidence and thirteenth in mortality among all cancers^1,2^. Although early-stage BC is typically treated with surgery, high rates of postoperative recurrence often require multimodal interventions^3,4^. For advanced or metastatic cases, systemic treatments are the current research hotspots, especially for immunotherapy^5^. Despite significant progress in recent years, 5-year recurrence-free survival rate still falls below 43%^6,7^. Currently, personalized precision therapy is gradually becoming the mainstream treatment. Immune checkpoint blockade (ICB) can elicit long-lasting responses in partial metastatic cancer patients^8^. For locally advanced or metastatic BC patients, who are refractory to platinum-based therapy, ICB has been regarded as a first-line or second-line treatment option^9^. Despite its great potential, only a small percentage of patients benefit from ICB (< 30%)^10,11^. The exact mechanisms and predictive factors that affect ICB efficacy remain unclear. Previous studies have identified some factors associated with ICB response, such as tumor immune microenvironment (TIME) patterns^12-14^, tumor mutational burden (TMB) and neoantigen load^15^, and microsatellite instability (MSI)^16^. These findings are essential for understanding the factors related to ICB response and developing predictive biomarkers. The current key obstacle for accurate therapy prediction is the search of ICB response biomarkers and resistance regulators^17,18^. Tumor molecular subtyping is a research trend for precision diagnosis and treatment, but currently limited subtyping method can accurately guide ICB therapy in BC patients^19^.

Within the tumor microenvironment (TME), tumor cells occupy specialized niches where they interact extensively with various factors, such as stromal and immune cells. These interactions have a significant impact on tumor initiation, progression, metastasis, and therapy response^20,21^. Studies have found that some cancer cell subsets can affect patients’ responses to immunotherapy, which are closely in contact with cancer-associated fibroblasts (CAFs) and CD8^+^ T cells (CD8Ts)^22,23^. Moreover, inhibitory cells, cytokines and metabolites that generate an immunosuppressive environment within the TME can reduce the activation and function of cytotoxic T-cells (CTLs), which will result in the tumor immune dysfunction (TID) or the exclusion of T-cells from the tumor (TIE)^12,21,24^. These two tumor-immune escape mechanisms will undermine tumor response to ICB therapy^17^. Liu et al. used tumor expression profiling data of melanoma and non-small cell lung cancer (NSCLC) to score these mechanisms and developed a computational framework called the Tumor Immune Dysfunction and Exclusion (TIDE) algorithm^17^. However, its effectiveness in BC, and further in pan-cancers, requires authoritative validation.

In this study, transcriptome analysis was conducted in the BC patients to assess TIDE status and identify specific biomarkers. Based on the identified marker-genes, a novel TIDE subtyping strategy was constructed which can classify BC patients into three subtypes with different clinicopathological and molecular features, prognoses, functional annotations, TME and therapy responses. This TIDE-based subtyping was also proven to accurately predict ICB treatment and chemotherapy outcomes in the BC patients. Furthermore, we validated the existence of the TIDE subtypes and their associations with ICB responses in five pan-cancer cohorts, which potentially indicates the consistency of our TIDE method across pan-tumors. Overall, we aimed to employ transcriptome data to evaluate TIDE status and develop an accurate TIDE-based decision-making tool in the BC and pan-cancer patients, which can facilitate precise treatment for tumors and gains valuable insights into the molecular mechanisms underlying tumor immune dysfunction and exclusion.

## Results

### 1 Correlations of TIDE status with the clinicopathological and molecular features in the BC patients

#### 1.1 Associations of TIDE scores with clinicopathological and molecular features

The workflow of our study is illustrated in **Figure S1** and **Supplementary Methods**. We utilized the TIDE algorithm^17^ to evaluate TIDE status and calculate TIDE scores based on bulk RNA-seq data. The samples were sorted from low to high to explore how TIDE scores were related to the clinicopathological and molecular features (**Figure 1A**). The results showed that younger patients, Asian ethnicity, and male gender had significantly lower TIDE scores (**Figure S2A**). TIDE scores were dramatically higher in those who died of BC, but unrelated to tumor progression (**Figure 1B, Figure S2B**). Additionally, patients with advanced pathological stage (T3 and T4), distant metastasis and high histological grade had higher TIDE scores, but not lymph node metastasis (**Figures 1C-1F**). Finally, TIDE scores of Papillary and Luminal papillary subtypes were the lowest among all histological subtypes and TCGA subtypes (**Figures 1G, 1H**).

**Figure 1.**
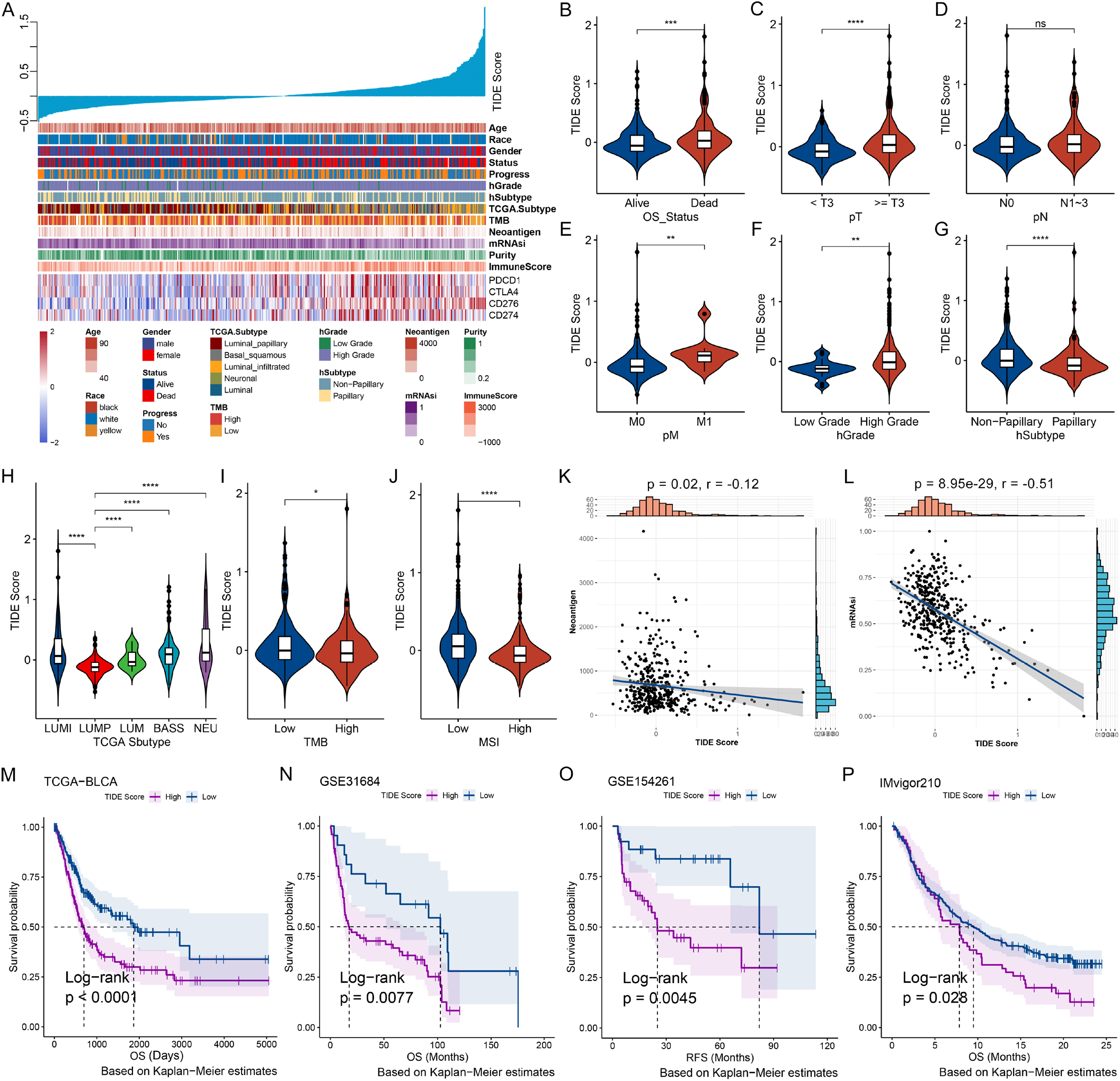
Correlations of TIDE status with clinicopathological and molecular features in the BC patients. **(A)** Associations of TIDE scores with clinicopathological and molecular features in BC patients. Columns represented samples ranked by TIDE scores from low to high (**top row**), and rows represent clinicopathological and molecular features associated with TIDE scores. **(B-J)** Comparisons of TIDE scores with subgroups of survival status (OS_Status) **(B)**, pathological TNM stage (pTNM) **(C-E)**, histological grade (hGrage) **(F)**, histological subtype (hSubtype) **(G)**, TCGA mRNA subtype **(H)**, tumor mutation load (TMB) **(I)** and microsatellite instability (MSI) **(J)**. LUMI, Luminal-infiltrated; LUMP, Luminal-papillary; LUM, Luminal; BASS, Basal-squamous; NEU, Neuronal. **(K, L)** Correlations of TIDE scores with neoantigen load **(K)** and stemness index (mRNAsi) **(L)** in BC patients. **(M-P)** Kaplan-Meier (K-M) analysis demonstrated a correlation of the TIDE scores with the prognosis of BC patients from TCGA-BLCA **(M)**, GSE31684 **(N)**, GSE154261 **(O)** and IMvigor210 **(P)** datasets. OS, overall survival; RFS, recurrence-free survival. Dashed line: median survival time. Color range: 95% confidence interval (CI). *p < 0.05, **p < 0.01, ***p < 0.001, ****p < 0.0001; ns, no significance.

Based on the somatic mutation data, we revealed that TIDE scores showed obviously negative correlations with TMB, MSI, neoantigen load, and stemness scores (mRNAsi) (**Figures 1I-1L**). For common biomarker mutation events, there were significant differences of TIDE scores between KDM6A, FGFR3, and TAF11 mutant patients and wild-type patients (**Figure S2C**).

Kaplan-Meier (K-M) analysis revealed significant associations of higher TIDE group with poorer overall survival (OS, **Figure 1M**), progression-free interval (PFI), and disease-specific survival (DSS) (**Figure S2D**). Additional five BC datasets confirmed these findings (**Figures 1N-1P, Figure S2E**). Therefore, based on the survival analysis results, tumor immune dysfunction and exclusion levels are probably two risk factors for BC.

#### 1.2 Correlation between TIDE status and TME

To determine the immune patterns of 431 TCGA_BLCA samples, we employed the ssGSEA algorithm^25^ to quantify scores for the 54 immune signatures published by Charoentong et al.^26^ and Şenbabaoğlu et al.^27^ The samples were divided into two groups: "High immune infiltration" (High-immu; n = 230, 53.4%) and "Low immune infiltration" (Low-immu; n = 201, 46.6%) (**Figure S3A**). Using the ESTIMATE algorithm to evaluate TME, the immune and stromal scores of High-immu group were significantly higher than the scores of Low-immu group (**Figure S3B**). DECEPTICON was applied to evaluate the immunocyte abundance. The results indicated that all types of immunocytes were significantly enriched in the High-immu group (**Figure S3C**), and a significant association was observed between the High-immu group and high TIDE scores, and vice versa (**Figure S3D**). Moreover, TIDE scores were positively correlated with Immune and Stromal scores, and negatively correlated with tumor purity and DNA fraction (**Figure S3E**). The myeloid cell infiltration was positively correlated with TIDE scores using Mantel’s test (**Figure S3E**).

### 2 Identification of three TIDE subtypes with significant differences in prognosis

#### 2.1 Identification of TIDE marker genes

Due to the correlation with TIDE scores and the prognosis of BC patients, we hypothesized that certain markers reflecting TIDE status could be used for molecular subtyping. Eight BC bulk RNA-seq datasets were applied to develop TIDE marker-genes. The detailed process is shown in **Figure S4A** and **Supplementary Methods**. Sixty-nine genes representing significant association with OS were selected (**Figure 2A**). These genes were classified into two clusters (**Figure S4B**). C1 comprises 11 genes that have no interaction with each other, while C2 includes 58 genes that were significantly enriched in signal pathways related to collagen and extracellular matrix metabolism, cell adhesion and integrin-mediated signal transduction, and epithelial-mesenchymal transition (**Figure 2A, Figure S4C**).

**Figure 2.**
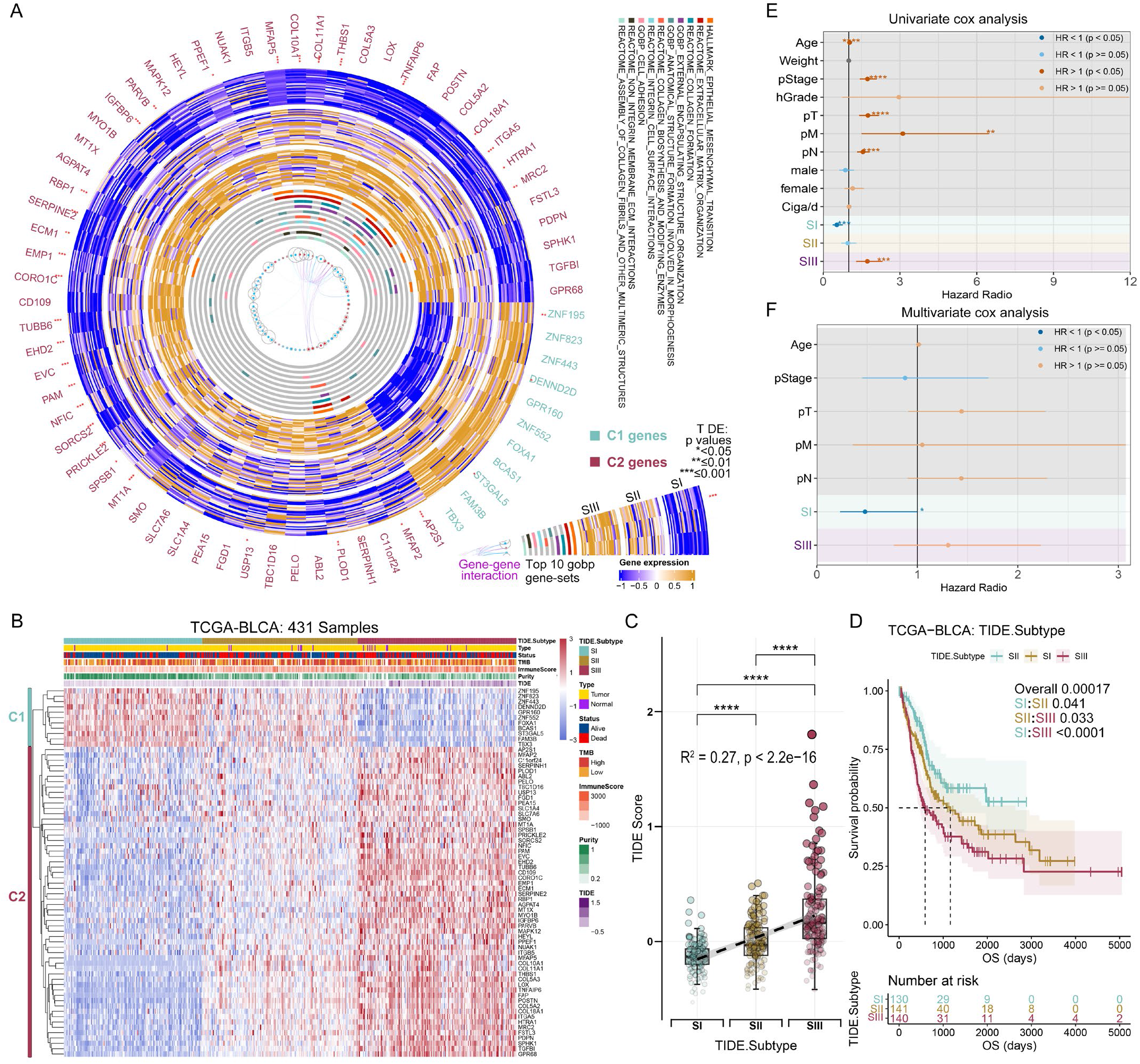
Identification of three TIDE-based subtypes of BC based on the TIDE marker genes. **(A)** CircosPlot shows the expression levels in TCGA-BLCA, signaling pathways and protein-protein interaction (PPI) networks of 69 TIDE marker genes. These genes are mainly divided into two clusters: C1 consists of 11 genes (cyan) and C2 comprises 58 genes (red). **(B)** Consensus clustering based on the expression of 69 TIDE marker genes classified patients of TCGA-BLCA into three subtypes: Subtype I (SI), Subtype II (SII), and Subtype III (SIII). C1 genes are labelled in cyan, and C2 are labelled in red. **(C)** Levels and trends of TIDE scores among three TIDE subtypes. **(D)** OS of TCGA-BLCA is significantly different among three TIDE subtypes. **(E, F)** Univariate **(E)** and multivariate **(F)** Cox regression analysis of the three TIDE subtypes with clinical and molecular characteristics. *p < 0.05, ***p < 0.001, ****p < 0.0001.

#### 2.2 Identification of three TIDE subtypes

Unsupervised consensus clustering^28^ was used to identify TIDE subtypes based on the expression profiling of 69 marker-genes (**Figure 2B**). Three clusters were well optimized determined with the help of consensus heatmap, cumulative distribution function (CDF) curves and proportion of ambiguous clustering algorithm (PAC) (35) (**Figure S5A, Figure 2B**). Subtype I (SI) consisted of 132 samples (30.6%) with high-expression of C1 genes and low-expression of C2 genes, subtype II (SII) comprised 148 samples (34.3%) with moderate-expression of both C1 and C2 genes, and subtype III (SIII) contained 151 samples (35.0%) with low-expression of C1 genes and high-expression of C2 genes (**Figure 2B**). The TIDE, Dysfunction and Exclusion scores were gradually increased from SI to SIII (**Figure 2C, Figure S5B**).

#### 2.3 Differences of prognosis among the TIDE subtypes in the BC patients

Based on K-M analysis, SIII represented the poorest prognosis, whereas SI owned the best prognosis, and SII showed an intermediate prognosis (**Figure 2D**, **Figure S5C**). By univariate Cox analysis, SI and SIII were protective and risk factors for OS, respectively (**Figure 2E**), and SI was an independent protective factor for OS by multivariate Cox analysis (**Figure 2F**).

#### 2.4 Stability and universality of TIDE subtyping strategy

Non-negative matrix factorization (NMF)^29^ and unsupervised hierarchical clustering were used to classify TCGA-BLCA patients (**Figures S5D-S5G**). The results were comparable to those obtained by using the consensus clustering. Therefore, TIDE subtyping strategy is robust across different algorithms. To further validate its universality, our novel TIDE subtyping strategy was also utilized in the other two integrated bulk RNA-seq cohorts. Both cohorts were consistently classified into three subtypes (**Figures S3H, S3I**) with significant TIDE scores and prognosis differences (**Figures S3J-S3M**). Therefore, these results indicate a good universality of the TIDE subtyping method.

### 3 Distinct clinical features, mutational events, and functional annotations among the TIDE subtypes

#### 3.1 Differences of clinicopathological features among the TIDE subtypes

The proportion of cancer adjacent samples were gradually increased from SI to SIII (**Figures 3A, 3B, Figure S6A**). TIDE subtypes have obvious differences in diagnostic age (**Figure 3C**), race and gender, but not in weight and daily smoking (**Figure S6A**). Regarding histopathology, SI had the lowest proportion of pT3 and pT4, high-grade (**Figures 3D, 3E**) and lymph node-positive patients (**Figure S6B**). We also observed a decreased proportion of patients with the Papillary histological subtype from SI to SIII (**Figure 3F**), and increased proportions of overall death, BC-specific death and progression **(Figure S6C)**.

**Figure 3.**
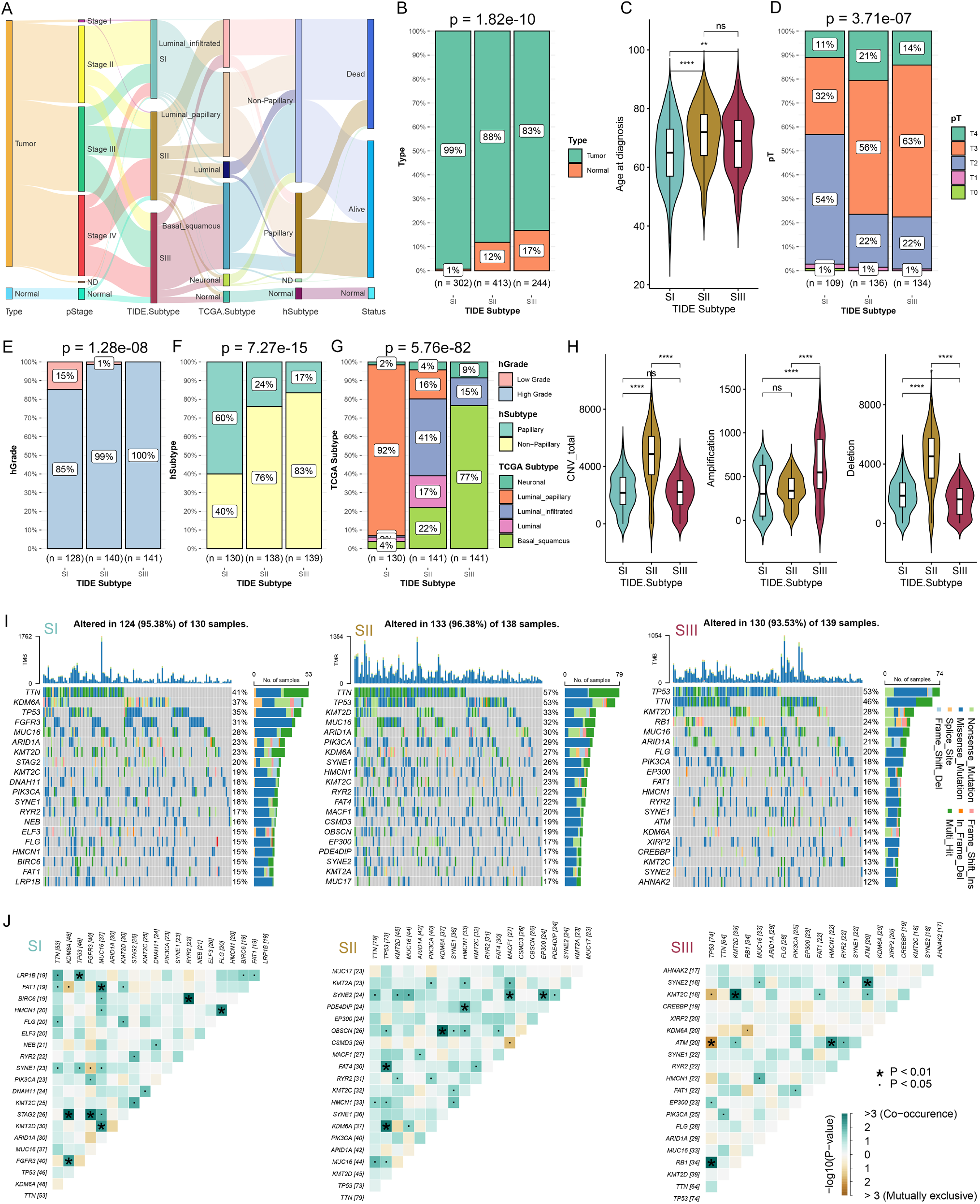
Comparisons of clinicopathological and molecular features among three TIDE subtypes in the BC patients. **(A)** Sankey diagram showing sample flows for TIDE subtype, sample type, pathological stage (pStage), TCGA Subtype, hSubtype, and survival status. **(B)** The proportion of sample type among the TIDE subtypes of UC.Combi cohort. **(C)** Comparisons of age at diagnosis among TIDE subtypes. **(D-G)** The proportion of pathological T stage (pT) **(D)**, hGrade **(E)**, hSubtype **(F)** and TCGA Subtype **(G)** among the three TIDE subtypes. **(H)** Comparisons of total copy number variations (CNV), amplifications and deletions among three TIDE subtypes. **(I)** Oncoplots showing the top 20 mutated genes in SI **(left)**, SII **(middle)**, and SIII **(right)**. **(J)** Heatmap showing mutually exclusive or co-occurrence events among top mutated genes in SI **(left)**, SII **(middle)**, and SIII **(right)**. *p < 0.05, **p < 0.01, ***p < 0.001, ****p < 0.0001; ns, no significance.

#### 3.2 Differences of molecular features among the TIDE subtypes

We further analyzed the compositions of different molecular subtypes in the TIDE subtypes and discovered obvious distribution differences, indicating a correlation and similarity between the known molecular subtypes and the new TIDE-based subtyping system (**Figure 3G, Figure S6D**). For example, the Luminal papillary subtype accounted for 92% in SI, and the Basal squamous subtype constituted 77% in SIII (**Figure 3G**). The SCNA and somatic mutation analysis showed that SII owned the highest copy number variation (CNV) burden (**Figure 3H**). However, SIII and SII displayed the highest amplification and deletion burdens, respectively. SII exhibited the highest level of TMB, as reflected by the total number of single nucleotide polymorphisms (**Figure S6E**). Each subtype showed the specific top mutated genes, with significant differences of mutation rates in the shared genes among the subtypes (**Figure 3I**). We examined gene co-mutation or mutual exclusivity patterns for each subtype (**Figure 3J**), which may affect BC subtyping, treatment, and prognosis. For instance, SIII carried co-mutated SYNE2 and ATM, two DNA damage repair genes. This rare co-mutation may increase DNA damage, tumor progression and metastasis, and alter tumor response to therapies^30^. Additionally, we investigated the mutation status of common BC biomarkers among the TIDE subtypes. TP53, PIK3CA, RB1, KDM6A, FGFR3ELF3, KMT2A, NFE2L2, FAT1 and SSH3 differed significantly in mutation proportions among subtypes (**Figure S6F**).

#### 3.3 Validation of the TIDE subtypes using the collected real-world BC samples

Thirty-one BC samples were collected for RNA sequencing to confirm the TIDE subtypes (**Figures 4A, 4B**). The detailed clinical information is shown in **Table S2**. The results indicated that the proportion of high-grade BC were increased from SI to SIII (**Figure 4C**). Among the 31 samples, the highest proportion of Ta was in SI and the highest proportion of T1 was in SII. T2, T4, and Tis stages belonged to SIII (**Figure 4D**). In terms of pathological diagnosis, SI possessed the highest proportion of papillary urothelial carcinoma, while SIII was consisted of the highest proportion of invasive urothelial carcinoma and urothelial carcinoma in situ (**Figure 4E**). Immunohistochemistry was used to assess the BC markers. The proportions of Ki67-positive cells, CCK5/6- and PHH3-positive samples, were the highest in SIII (**Figures 4F-4H**), indicating a close association with BC malignancy and invasiveness. CD44 signal was the lowest in SIII subtype (**Figure 4I**). The published data suggested that loss of CD44 staining may indicate defective urothelial development or urothelial carcinoma in situ. Finally, we found significant differences in tumor volume among the subtypes based on CT imaging data (**Figure 4J**). Tumors belonging to SI were generally smaller, while those belonging to SIII were often larger or even infiltrated the entire bladder (**Figure 4K**). These results consistently demonstrate that the SIII subtype has the highest malignancy and is closely associated with urothelial carcinoma in situ.

**Figure 4.**
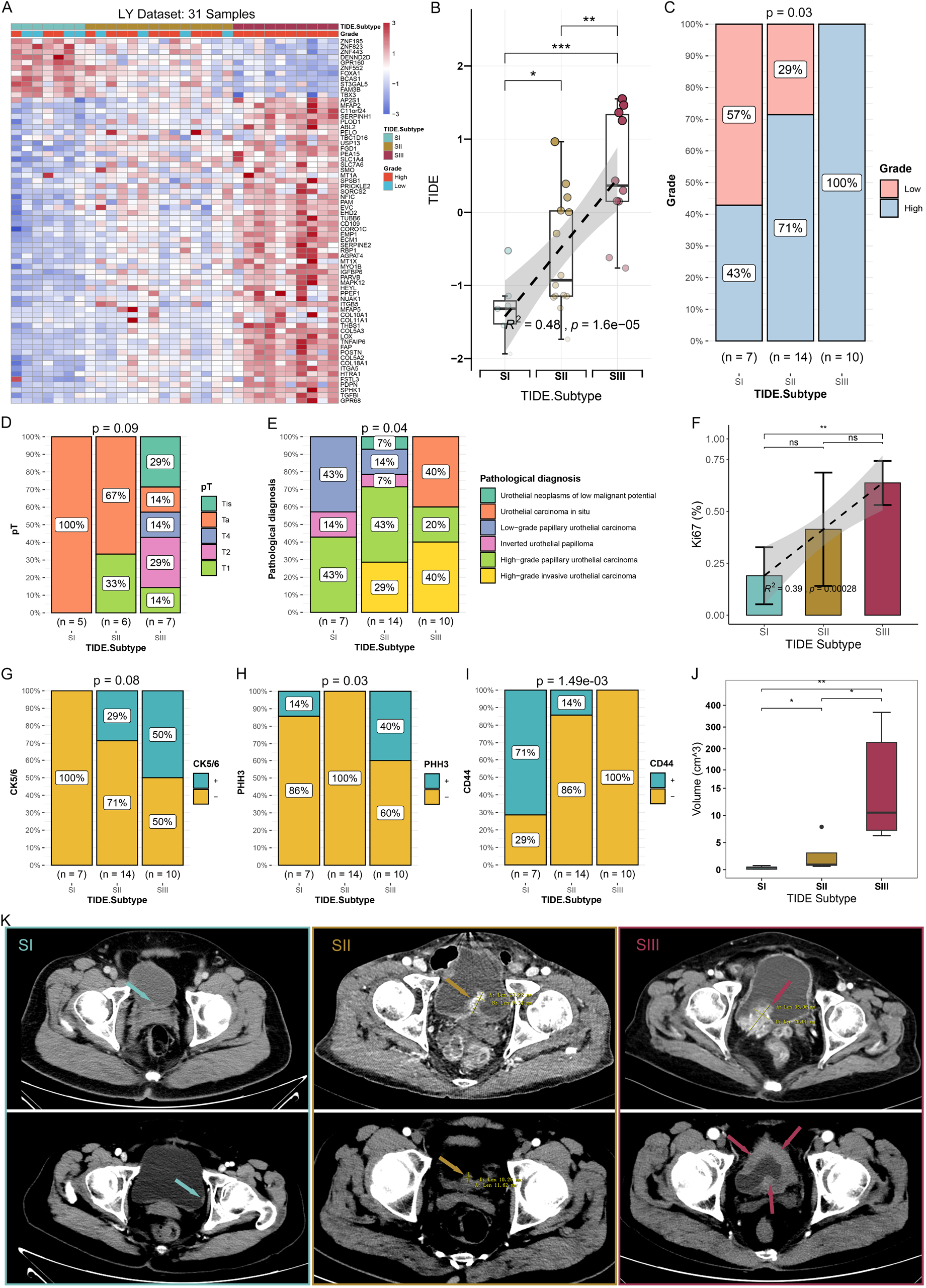
Validation of the TIDE subtypes using the collected real-world BC samples. **(A)** Consensus clustering by the TIDE marker genes can distinguish the real-world BC samples into three TIDE subtypes. **(B)** Levels and trends of TIDE scores among three TIDE subtypes. **(C-E)** The proportion of the tumor grade **(C)**, pT **(D)** and pathological diagnosis **(E)** among the TIDE subtypes. **(F)** Percentage of Ki67 positive cells in three TIDE subtypes. **(G-I)** The proportions of positive CK5/6 **(G)**, PHH3 **(H)** and CD44 **(I)** samples in three TIDE subtypes. **(J)** Tumor volumes of three TIDE subtypes based on CT tomography. **(K)** Typical CT tomographic images of bladder tumors (→) in three TIDE subtypes. *p < 0.05, **p < 0.01, ***p < 0.001. ns, no significance.

#### 3.4 Distinct signaling pathways and functional annotations among the three TIDE subtypes

Gene Set Variation Analysis (GSVA), Gene Set Enrichment Analysis (GSEA), and Ingenuity Pathway Analysis (IPA) were conducted to investigate the functional annotations and potential mechanisms associated with TIDE subtypes. GSEA showed SI had enriched signaling pathways related to lipid and xenobiotic metabolism (**Figure S7A**). In contrast, SII displayed enriched pathways related to complement activation, humoral immunity, scavenger receptor and B cell receptor (**Figure S7B**). SIII showed enriched pathways related to various cellular processes, including migration, activation, development, proliferation, inflammation and immune regulation (**Figure S7C**). These results were consistent with the observation from GSVA (**Figure S7D**). IPA analysis of canonical signaling pathways were used to reveal the status of activation or inhibition. Our results showed that SI activated biosynthesis and xenobiotic metabolism pathways, but suppressed cancer, stress, cytokine, immune and growth pathways **(Figure S7E)**. SIII displayed opposite pathway activity to SI **(Figure S7I)**. SII was a transitional state with mixed pathway signals **(Figure S7G)**. The graphical summary suggested that SI suppressed immune and inflammatory factors, while SIII activated them (**Figures S7F, S7J, S7K**). Lastly, consistent with GSVA and GSEA, SI activated lipid metabolism pathways, but SIII suppressed them (**Figure S7L**).

### 4 TME patterns of three TIDE subtypes

TME is a highly researched topic, as it plays a crucial role in the initiation, advancement, and management of tumors, specifically in ICB treatment^31^. Therefore, we attempted to characterize the TME patterns for TIDE subtypes. We observed a significant decrease in tumor purity and DNA fraction, and a significant increase in ESTIMATE, Stromal and Immune scores, and the proportion of High-immu subtype from SI to SIII (**Figure 5A, Figure S8A**). Hypergeometric test indicated that SI was significantly associated with Low-immu subtype, but the SII and SIII subtypes were significantly associated with the High-immu subtype (**Figure 5B**). Additionally, we examined the association of TIDE subtypes with immune phenotypes. It displayed that the desert phenotype was mainly in SI, the inflamed phenotype was primarily in SIII, and the excluded phenotype was mainly in SII (**Figure 5C**). These results indicated that the proportion of immune and stromal cells was increased gradually from SI to SIII. Using the SCDC algorithm^32^, we calculated the immunocyte abundance in TCGA-BLCA patients. A gradual increase in myeloid cells, CD8Ts and CAFs, and a decrease in B cells, CD4Ts and endothelial cells was found from SI to SIII (**Figure 5D**). To further validate our findings, the DECEPTICON and TIDE algorithms^17^ were used to evaluate TCGA-BLCA and IMvigor210. The results were consistent with our findings (**Figures S8B-S8E**). H&E staining of the real-world BC samples showed the gene expression patterns of the TIDE subtypes were histologically related to the abundance of stromal and tumor cells. SI represented abundant tumor cells, and SIII showed highly fibrotic stroma with abundant immunocytes **(Figure 5E)**. These pathological features were verified using diagnostic slides of TCGA-BLCA **(Figures S8F)**.

**Figure 5.**
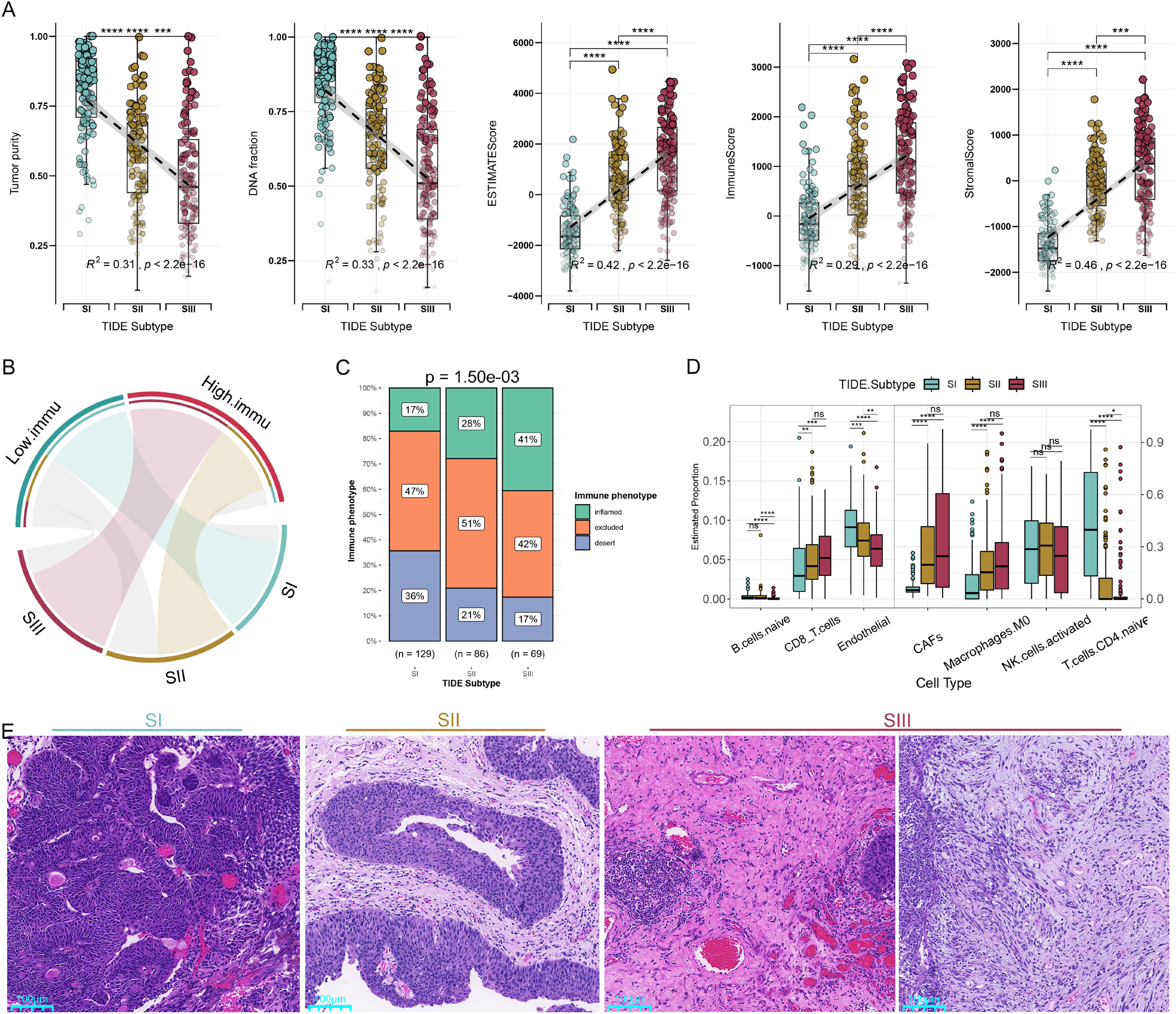
Characterization of TME patterns in the TIDE subtypes based on bulk RNA-seq datasets. **(A)** Differences of tumor purity, DNA fraction, and TME scores among the TIDE subtypes. **(B)** Hypergeometric tests revealed an association between TME patterns and TIDE subtypes. Gray lines represent no significance. **(C)** The proportion of immune phenotype among the TIDE subtypes of TCGA-BLCA. **(D)** Boxplots showing comparisons of stromal cells abundance among the three TIDE subtypes of TCGA-BLCA. Cell proportions are assessed by the SCDC algorithm. **(E,)** Representative H&E histological images of the three subtypes from the real-world BC samples. *p < 0.05, **p < 0.01, ***p < 0.001, ****p < 0.0001; ns, no significance; scale bar, 100 μm.

Subsequently, a scRNA-seq dataset was further analyzed to describe the immune landscape of TIDE subtypes. Single-cells were visualized via Uniform Manifold Approximation and Projection (UMAP), and were divided into 31 clusters (**Figure S9A**). Each cluster was annotated as a specific cell-type (**Figure 6A, Figure S9B**). The expressions of TIDE marker-genes were examined in each cell-type and found that C1 genes were mainly expressed in tumor cells, and C2 genes were highly expressed in monocytes, macrophages, fibroblasts, and endothelial cells (**Figure 6B**). The results confirmed that SI enriched of C1 genes brought higher tumor purity, while SIII enriched of C2 genes carried higher immune and stromal components. To obtain the corresponding TIDE subtypes of these patients, pseudobulk data was obtained from each patient based on scRNA-seq data and consensus clustering^28^ was conducted. Patients were divided into two subtypes (**Figure S12D**): low expression of C1 and C2 genes (Low), and high expression of C1 and C2 genes (High). By comparing the cell composition, we found that the proportions of monocytes, macrophages, fibroblasts, CD8Ts and endothelial cells were higher in the High subtype, whereas the proportions of Tregs, B cells and plasma cells were at the lower abundance (**Figure 6C**). This finding is consistent with our previous deconvolution results. Next, the terminally exhausted signals^33^ were evaluated and indicated the High subtype of CD8Ts exhibited a higher level of terminal exhaustion status (**Figure 6D**). By trajectory analysis, the High subtype of CD8Ts differentiated into two branches was found as the tumor progressed (**Figures 8E, 8F**). The late differentiating group of CD8Ts did not exhibit significant changes in exhaustion status, whereas the early group displayed severe exhaustion **(Figures 8E, 8G**). This specific branch of CD8Ts may represent a critical population for poor prognosis and ICB resistance in High subtype patients. In contrast, the exhaustion status decreased as the tumor progressed in the Low subtype (**Figures 8E, 8G**). Overall, the trend of C2 expression on the trajectory was consistent with the terminal exhaustion signature and further affirmed the reliable subtyping by C2 genes (**Figure S9E**).

**Figure 6.**
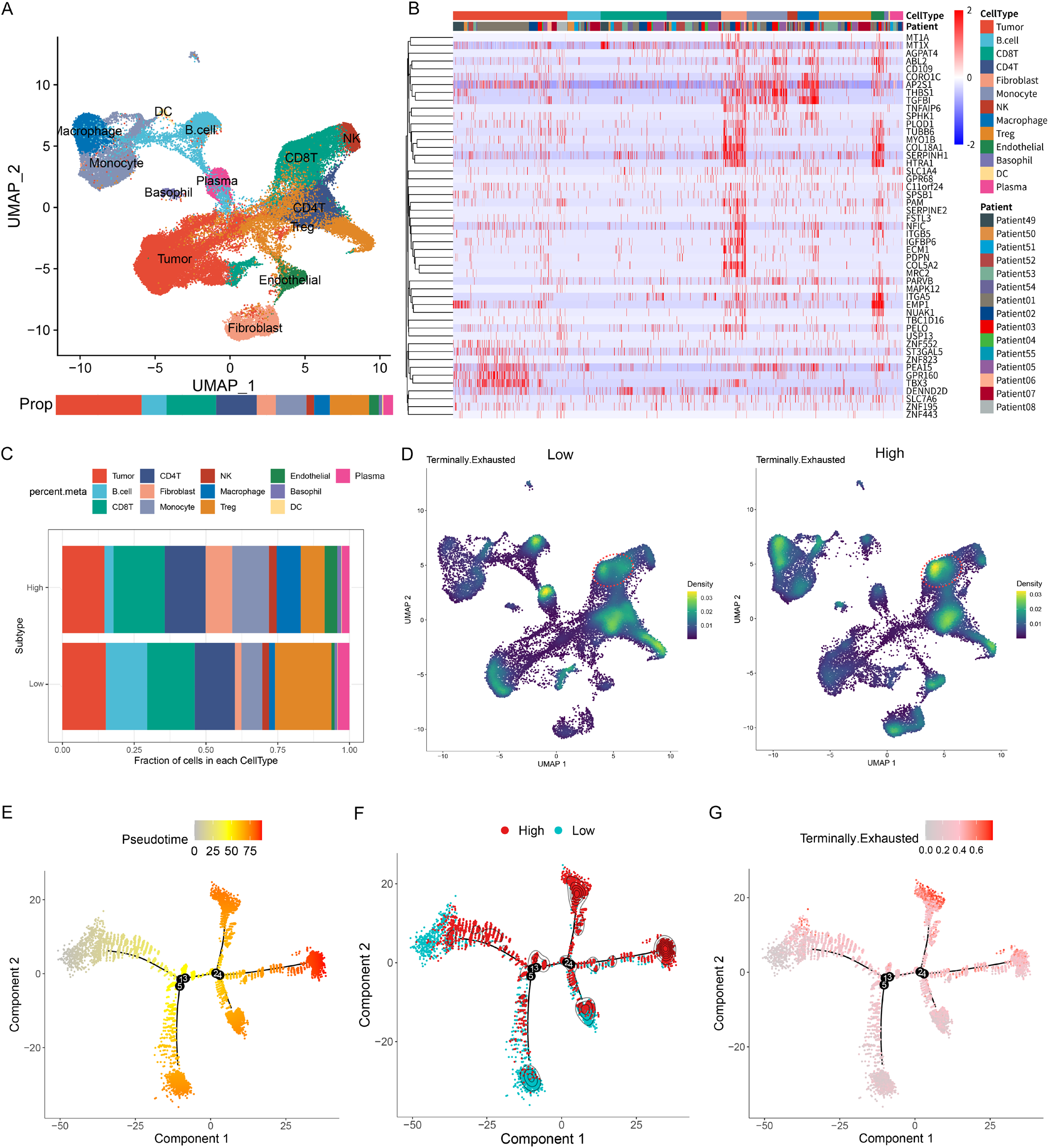
Characterization of TME patterns in the TIDE subtypes based on single-cell RNA-seq dataset. **(A)** Uniform Manifold Approximation and Projection (UMAP) plot was used to analyze the single-cell RNA-seq dataset. Each color represents one of the 13 cell types in Salomé’s dataset. **(B)** Heatmap showing the expression levels of TIDE marker genes in 13 cell types. **(C)** Proportions of 13 cell types in TIDE subtypes. **(D)** Terminally exhausted signals of two subtype cells assessed by weighted kernel density estimation. Red circle marks CD8Ts. **(E, F)** Differentiation trajectory of CD8^+^ T cells (CD8Ts) in BC, with a color code for pseudotime **(E)** and TIDE subtypes **(F)**. **(G)** Expression levels of terminally exhausted signature along the CD8Ts differentiation trajectory in BC.

### 5 Precision treatment of BC following the TIDE subtypes

#### 5.1 Differential expression of immune checkpoint molecules among the TIDE subtypes

Since tumor molecular subtyping is pivotal to provide a basis for precise diagnosis and personalized treatment, we further evaluated the potential of TIDE subtypes in guiding precision treatment and drug selection. We found that TIDE score was positively correlated with most immune checkpoint molecules and ligands, except for a few molecules such as CEACAM1 and TNFRSF25 (**Figure S10A**). Notably, most immune checkpoint molecules and ligands (such as PD-1, PD-L1, CTLA4, etc.) were increased from SI to SIII (**Figure 7A, Figure S10B**). In the TIME, CD8Ts are often suppressed and exhausted by immune checkpoint molecules. Meanwhile, exhausted CD8Ts typically express high levels of immune checkpoint molecules^34^. In our study, SIII patients were possibly under this exhausted status, and the observed TIME patterns along with the differential expressions of immune checkpoint molecules could be critical for therapeutic efficacy.

**Figure 7.**
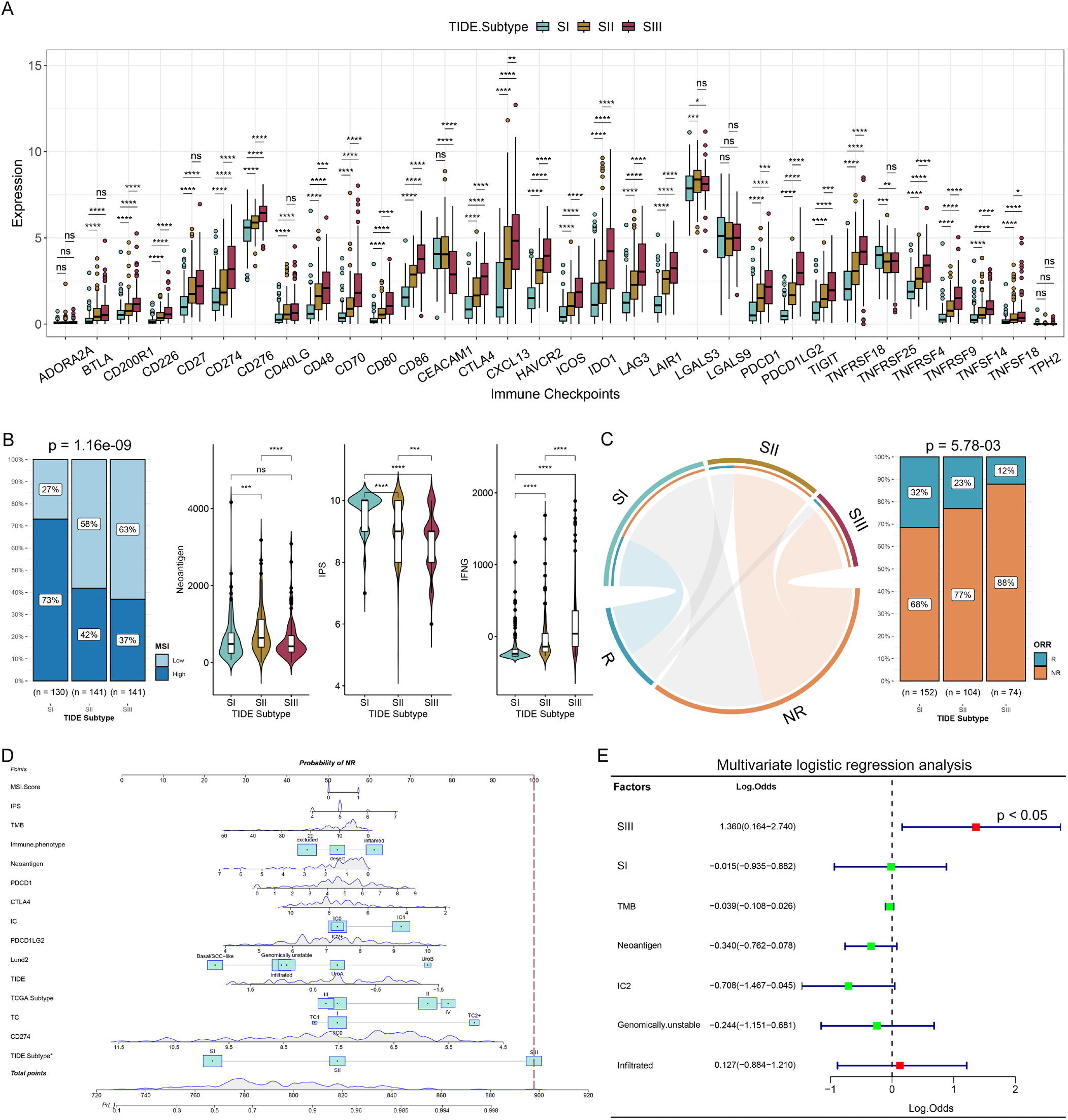
TIDE subtypes were closely related to ICB responses. **(A)** Comparisons of expression levels of immune checkpoint molecules among the TIDE subtypes of TCGA-BLCA. **(B)** Comparisons of MSI, neoantigen load, IPS and interferon gamma among the TIDE subtypes of TCGA-BLCA. **(C)** Hypergeometric test revealed an association between TIDE subtypes of ICB.UC cohort and ICB responses **(left)**, gray lines represent no significance; Stacked histogram showing the differences of ICB responses among the TIDE subtypes of ICB.UC cohort **(right)**. **(D)** Nomogram for predicting ICB non-responders based on logistic regression model. Red line showing the SIII points. **(E)** Impacts of the TIDE subtypes and other predictive biomarkers on ICB efficacy, which were achieved by multivariate logistic regression analysis. R, response; NR, non-response. *p < 0.05, **p < 0.01, ***p < 0.001, ****p < 0.0001; ns, no significance.

#### 5.2 Distinct sensitivity to ICB among the TIDE subtypes

We initially examined the profile of the predictive ICB biomarkers in TIDE subtypes. The results showed that the proportions of IC1 and IC2+, as well as TC1 and TC2+, were significantly higher in SIII (**Figure S10C**). IC and TC can indicate the PD-L1 levels of immunocytes and tumor cells in the TME patterns. The interferon gamma (IFNG) score was gradually increased from SI to SIII (**Figure 7B**). Conversely, the proportion of high MSI patients, neoantigen load and immune phenotype score (IPS) were significantly lower in SIII. (**Figure 7B, Figure S10D**). Notably, ICB response tends to increase along with the higher level of the abovementioned biomarkers^16,18,26,35^. Therefore, we can find that the prediction of the known ICB biomarkers contains conflicts in evaluating the ICB efficacy of TIDE subtypes.

We further analyzed the ICB response rates of TIDE subtypes in the ICB.UC cohort. The results indicated that SII and SIII were significantly associated with non-responders (NR), SI was closely associated with responders (R), and the proportion of responders were gradually decreased from SI to SIII (**Figure 7C**). The analysis of the IMvigor210 dataset produced the consistent results (**Figure S10E**). We also analyzed known biomarkers in predicting ICB responses using the IMvigor210 dataset (**Figure S10F**). We only found that the response rates between low and high TMB differed significantly. Nanogram showed the SIII contributed the most to ICB non-response (**Figure 7D**). Univariate logistic regression analysis indicated that SI was a protective factor, but SIII was a risk factor for ICB response (**Figure S10G**). The multivariate logistic regression analysis showed that SIII was the most significant independent predictor of non-response (**Figure 7E**, log odds = 1.36, p = 0.036). Finally, we analyzed the predictive value of TIDE subtypes in combination with other biomarkers for ICB outcomes. Receiver operating characteristic (ROC) curves showed that the combined approach of TIDE subtypes and other biomarkers had significantly improved predictive values compared to a single biomarker (**Figure S12H**). These findings suggest that TIDE subtype can compensate, to some extent, for the limitations of using a single biomarker in predicting ICB response.

#### 5.3 TIDE subtypes retain sensitivity to specific drugs

We assessed the sensitivity of TIDE subtypes to targeted drugs. As shown in **Figure S11A**, the SIII subtype tended to be benefited from anti-vascular endothelial growth factor antibodies (anti-VEGF), heat shock protein inhibitors (HPSIs), and poly ADP ribose polymerase inhibitors (PARPIs). On the other hand, according to Fisher’s test results, SI patients tended to be resistant to anti-human epidermal growth factor receptor 2 antibodies (anti-HER2/ERBB2) and PARPIs, and the SII group tended to be resistant to mammalian rapamycin target protein inhibitors (mTORIs).

Subsequently, we evaluated the sensitivity of TIDE subtypes to the first-line treatments and recommended drugs. Our findings indicated that certain subtypes showed differential sensitivities to particular drugs. For example, among antimetabolites, SI showed higher sensitivity to methotrexate and gemcitabine (**Figures S11B, S11C**), while SIII was more sensitive to pemetrexed (**Figure S11D**). Among plant alkaloids, SI was more responsive to vincristine (**Figure S11E**), and SIII showed higher sensitivity to paclitaxel (**Figure S11F**). Additionally, SI demonstrated higher sensitivity to anti-tumor antibiotics, such as doxorubicin and epirubicin (**Figures S11G, S11H**), while SIII was more sensitive to bleomycin (**Figure S11I**). Furthermore, our analysis revealed that SIII exhibited higher sensitivity to platinum drugs, including cisplatin and oxaliplatin (**Figures S11J, S11K**), alkylating agents such as ifosfamide, mitoxantrone and fludarabine (**Figures S11L-S11N**). In terms of the second-line or recommended drugs, our analysis showed that SI was less responsive to fibroblast growth factor receptor (FGFR) inhibitors, such as erlotinib and AZD4547 (**Figures S11O, S11P**), as well as to the ERBB2 inhibitor DMOG_165 (**Figure S11Q**). On the other hand, SIII exhibited higher sensitivity to epidermal growth factor receptor inhibitors (EGFRIs), like gefitinib (**Figure S11R**), PARPIs (e.g., olaparib, rucaparib, talazoparib and niraparib) (**Figures S11S-S11V**), as well as to imatinib (**Figure S11W**), and the anti-EGFR antibody cetuximab (**Figure S11X**).

### 6 Validation of the TIDE-based subtyping strategy in pan-tumors

Pan-cancer research enables the application of diagnosis and treatment to a broader range of tumor types by identifying commonalities among them. To determine whether our TIDE subtypes were conserved in pan-cancer, we analyzed five pan-cancer datasets. We initially classified the TCGA pan-cancer samples into three subtypes using 69 TIDE marker genes (**Figure S12A-left**). We observed that 11 C1 genes were similarly expressed in pan-cancer samples, and the pan-cancer samples cannot be clustered using these genes (**Figure S12A-right**). Therefore, we excluded these 11 genes and used the remaining 58 C2 genes to classify the pan-cancer samples. The results showed that the five pan-cancer datasets were consistently classified into three subtypes: SI (low expression), SII (medium expression), and SIII (high expression) (**Figure 8A, Figures S12C-S12F**). Moreover, there were significant differences in prognosis among the subtypes (**Figure 8A, Figures S12B-S12D**), indicating that TIDE subtypes are ubiquitous in pan-tumors.

**Figure 8.**
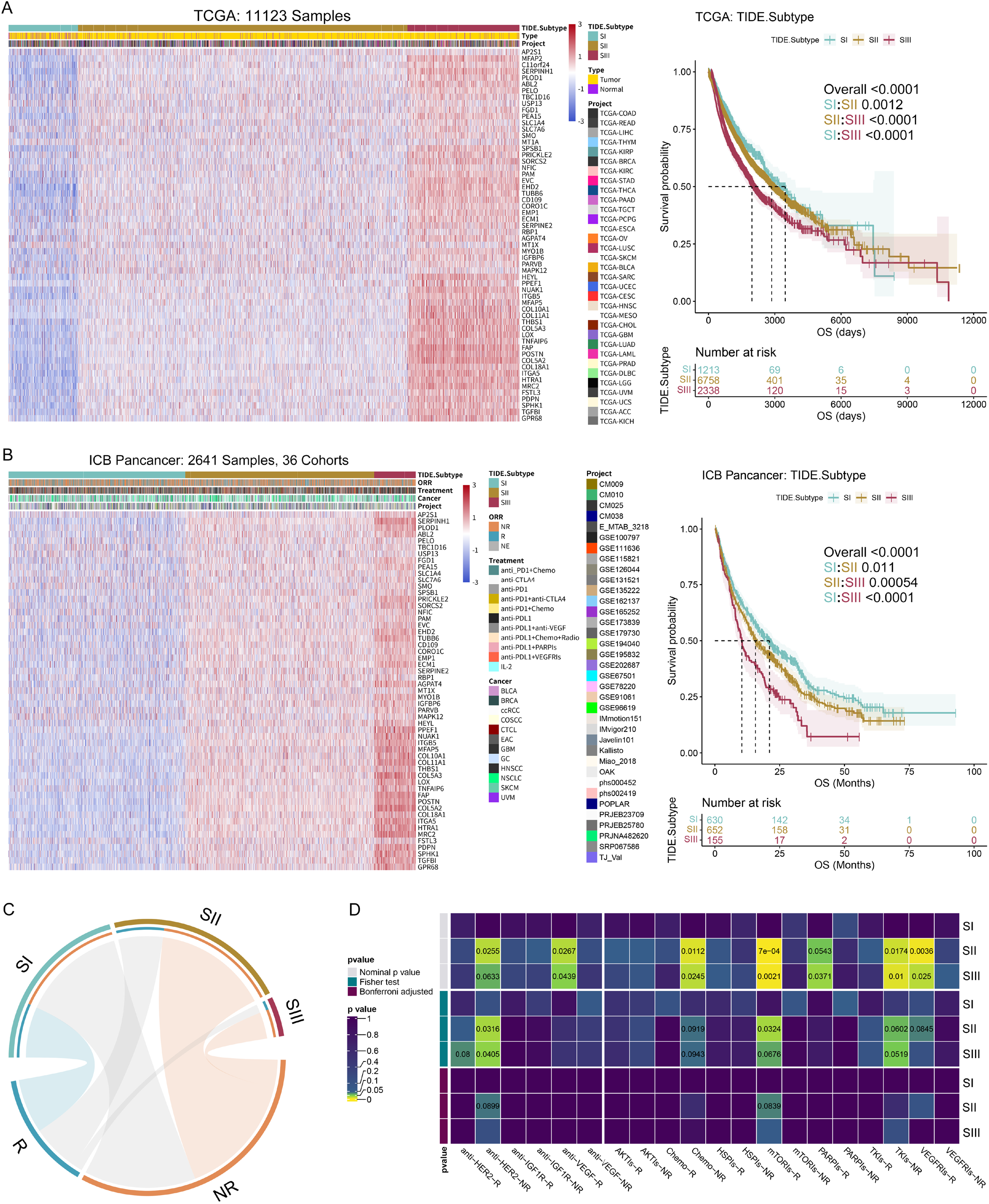
Conservations of the three TIDE subtypes in pan-tumors. **(A)** Unsupervised hierarchical clustering using the 58 C2 genes to classify pan-tumor samples into three subtypes from TCGA patients **(left)**. K-M analysis shows distinct OS of the TIDE subtypes **(right)**. **(B)** The same analysis as A. Unsupervised hierarchical clustering of Pan-cancer samples treated with ICB is shown on the left, and K-M analysis showing distinct OS of the TIDE subtypes is on the right. **(C)** Hypergeometric test collaborates an association of pan-tumor TIDE subtypes with ICB therapy responses. Gray lines represent no significance. **(D)** Submap analysis reflects the sensitivity of the pan-tumor TIDE subtypes to the targeted treatments.

ICB exerts a remarkable therapeutic effect for many tumors. So, we explored the sensitivity of TIDE subtypes to ICB treatment in pan-tumors. We obtained RNA-seq data from 2641 patients across 12 tumor types. Based on the expression levels of C2 genes, we assigned these samples into three subtypes with significantly different prognoses (**Figure 8B, Figure S12G**). Hypergeometric test showed that ICB responders were closely related to SI and non-responders were associated with SII and SIII (**Figure 8C**). Stacked bar displayed a significant decrease in the response rates from SI to SIII (**Figure S12H**). Submap analysis indicated that SIII was mainly correlated with anti-PD-L1 resistance, while SII was associated with IL-2 treatment response (**Figure S12I**). Regarding other targeted drugs, SII and SIII were responsive to anti-VEGF, mTORIs, PARPIs, VEGFRIs treatments, but were resistant to anti-HER2 and TKIs (**Figure 8D**).

## Discussion

Tumor immune dysfunction and exclusion are regarded as two main mechanisms of tumor immune escape. The former refers to T cell dysfunction in tumors with high cytotoxic T lymphocyte abundance^17,36^. The persistent antigen stimulation or immunosuppressive signals can trigger functional decline or exhaustion in CTLs, with lower cytokine production and co-stimulatory molecule expression, and higher expression of immune checkpoint receptors and immunosuppressive enzymes^37^. Tumor immune exclusion refers to preventing T cell infiltration in tumors with low CTL abundance, known as "cold tumors"^36^. Tumor cells utilize different mechanisms to obstruct T cell infiltration. The secretion of immunosuppressive factors, like TGF-β, IL-10, and IL-35, will hinder T cell activation and proliferation^38^. The expression of immune checkpoint molecules from cancer cells, that interact with the corresponding receptors and deliver negative signals, impede T cell activity or induce tolerance^34^. The alteration of the physical and metabolic TME characteristics also hamper T cell migration and function, such as interstitial fibrosis, lactate production and hypoxia^12,39^. The recruitment of other immunosuppressive cells that compete with T cells, such as Tregs, myeloid-derived suppressor cells (MDSCs), and tumor-associated macrophages, will ultimately suppress their anti-tumor response^21,24,39^.

The development of bioinformatics techniques has propelled unprecedented advances in precision medicine, such as genomic and transcriptomic analyses, and multi-omics integration analysis^40^. Elucidation of the specific treatment response changes in the TME patterns and immune cell functional status can provide insights into mechanisms underlying resistance of immune therapy and identify new therapeutic strategies. In our study, transcriptome data was used to evaluate the status of TIDE patterns, and a novel TIDE-based subtyping method was developed for accurately predicting immunotherapy in the BC patients. We found SIII subtype owning the highest immune infiltration. The cytokine, inflammatory response, and immune regulation pathways were highly enriched and activated. The TME analysis indicated that SIII represented the lowest tumor purity, but the highest amount of fibrosis and immunosuppressive cells (macrophages and MDSCs), that would impair ICB efficacy. Moreover, the SIII subtype exhibited elevated CTLs infiltrations, immune checkpoints and ligands, and exhausted CD8Ts, indicating functional defects and exclusion of CTLs in the TME. This immune dysfunction and exclusion would affect immunotherapy and chemotherapy^12,17^. Additionally, SIII represented reduced B cells and plasma cells, which would also undermine ICB efficacy^22,41,42^. Overall, the heterogeneity of TME reflects the resistance of SIII to ICB treatment in the BC patients.

Tumor molecular subtyping is pivotal for cancer diagnosis and treatment, which helps to reflect the TME patterns, select ICB candidates, predict immune-related adverse events, and guide ICB combination strategies^19,43^. The newly generated TIDE-based subtyping method was further compared with other known methods. Based on the immune phenotypes, SIII represented the highest inflamed phenotype, which normally indicates more responsive to ICB therapy due to high PD-L1 level and abundant tumor-infiltrating lymphocytes^44^. Moreover, it is believed that increased IC1 and IC2+ or TC1 and TC2+ indicate higher objective immunological remission rates^18^. SIII exhibited the highest proportion of IC and TC levels among the subtypes, but in fact, SIII failed to respond to ICB. Therefore, these results indicate the accurate prediction performance of TIDE-based subtyping method in identifying non-responders compared with certain immunotherapy biomarkers. The outcome was also confirmed by multivariate logistic regression. It might suggest an option to combine TIDE subtypes with biomarkers for guiding ICB treatment. We can select patients with inflamed phenotype tumors and further exclude SIII subtype patients for ICB therapy candidates. Based on the IMvigor210 cohort, ROC showed that the immune phenotype alone had lower predictive performance (AUC = 0.55), but it was notably improved with the combination of TIDE subtyping (AUC = 0.65). However, large-scale clinical trials are still needed for future validation.

The TIDE algorithm developed by Liu et al. integrates signatures of T cell dysfunction and T cell exclusion. It exhibited excellent predictive performance for ICB efficacy in melanoma and NSCLC with an AUC up to 80%^17^. However, there is still a lack of evidence to support this method to predict ICB response in BC. Our study showed that the original TIDE algorithm had limited predictive performance for ICB response, and no differences of the ICB response could be detected by the TIDE algorithm. ROC curve also indicated that the TIDE scores had little predictive power for ICB efficacy (AUC = 0.57). In our study, we identified three TIDE subtypes closely associated with ICB efficacy, which optimized the applicability of the algorithm in BC. We characterized the TIDE status of BC and developed a TIDE subtyping method. Briefly, SI represents the lowest TIDE status, the best prognosis, and it is sensitive to ICB treatment. SIII shows the highest TIDE level, the poorest prognosis, accompanied by suppressive TIME and terminally exhausted T cells, which is sensitive to EGFRIs and PARPIs but resistant to ICB treatment. SII falls into a transitional status with intermediate TIDE level and prognosis. We also validated the conserved characteristics of TIDE subtypes in pan-cancers using five pan-tumor cohorts, one ICB pre-treatment cohort and their responses to immunotherapy. These results suggest that our novel TIDE-based subtyping strategy can be also used for pan-cancers, and potentially bring more benefits for a wider range of cancer patients.

In conclusion, tumor molecular subtyping represents one future research direction for the personalized cancer therapy, with important implications in guiding clinical treatment and developing new anti-cancer drugs. Our analysis of whole transcriptome data in the BC patients identified three TIDE-based subtypes showing significant differences in clinicopathological and molecular features, as well as functional pathways and treatment responses. This subtyping method can also be applied to pan-cancer. We believe that our novel TIDE-based subtyping strategy has enormous potential for clinical application, as it can assist in making personalized treatment decisions for BC and pan-cancer patients, selecting potential beneficiaries, and excluding resistant patients of ICB therapy.

## Methods

Five bulk RNA-seq datasets and one scRNA-seq dataset of BC, five bulk-RNA-seq cohorts of pan-tumors, one bulk RNA-seq cohort of pan-tumors treated with ICB, and somatic mutation and CNA data from TCGA-BLCA were collected in this study. Details and sources for all datasets are listed in **Table S1**. The overall design of this study is as follows: evaluation of TIDE status and its relationship with BC clinicopathological and molecular features; TIDE subtyping; characterization of clinicopathological and molecular features of TIDE subtypes; and analysis of the pan-tumor landscape of the TIDE subtypes. The analysis methods used include RNA sequencing, DE analysis, clustering analysis, TIDE analysis, PPI, pathway analysis, somatic mutation and CNV analysis, survival analysis and immunohistochemistry, etc. Detailed descriptions of the methods and computational analyses are provided in **Supplementary Information**.

## Declarations

### Ethics approval and consent to participate

The institutional review board of Shanghai Sixth People’s Hospital approved this study, and informed consent was obtained from all patients. The ethical approval number is 2023-KY-131 (K).

### Consent for publication

Written informed consent for publication of their clinical details and/or clinical images was obtained from the patient/parent/guardian/ relative of the patient.

### Availability of data and material

The original contributions presented in the study are included in the article and supplementary information. The RNA-seq data of the collected BC in the LY dataset has been uploaded to GEO database. The computer R code for processing and analysis of this study is available upon request.

### Author’s Contributions

Conceptualization: K.N., X.H.; Methodology: K.Z., K.N., and X.H.; Investigation: K.Z., Y.H., and H.C.; Project administration: K.N. and X.H.; Writing original draft: K.Z., Y.H., H.C., X.H., and K.N. All authors have read and approved the final manuscript.

## Acknowledgements

The part computations in this study were run on the Siyuan-1 cluster, supported by the Center for High Performance Computing at Shanghai Jiao Tong University.

