## Supplementary Information for "Three Conserved Immune Dysfunction and Exclusion Subtypes in Bladder and Pan-cancers: Prognostic and Immunotherapeutic Significance"

#### Supplementary Tables


- Supplementary Table 1
- Supplementary Table 2


#### Supplementary Figures


- Supplementary Figure 1
- Supplementary Figure 2
- Supplementary Figure 3
- Supplementary Figure 4
- Supplementary Figure 5
- Supplementary Figure 6
- Supplementary Figure 7
- Supplementary Figure 8
- Supplementary Figure 9
- Supplementary Figure 10
- Supplementary Figure 11
- Supplementary Figure 12


#### Supplementary Methods

#### Supplementary References

**Supplementary Table 1.** Characteristics of single-cell RNA-seq dataset and bulk RNA-seq datasets enrolled in this study.

**a.** Bulk RNA-seq datasets of bladder cancer

| Datasets | Samples | | Age (yr) | pT/cT | | | | | pN/cN | | pM/cM | | Gender | | Survival | TMB | Purity | Ref. | Download |
|---|---|---|---|---|---|---|---|---|---|---|---|---|---|---|---|---|---|---|---|
| | T | N | | T0 | T1 | T2 | T3 | T4 | N0 | N1-N3 | M0 | M1 | Male | Female | | | | | |
| **TCGA-BLCA** | 412 | 19 | 69 (34-90) | 1 | 5 | 123 | 207 | 62 | 249 | 138 | 211 | 11 | 314 | 117 | OS, PFI, DSS, DFI | √ | √ | | TCGA |
| **GSE31684** | 93 | | 69 (42-91) | 5 | 10 | 17 | 42 | 19 | 49 | 28 | 57 | 36 | 68 | 25 | OS, DSS, RFS | | | [1] | GEO |
| **GSE154261** | 99 | | | | | | | | | | | | | | RFS, PFS | | | [2] | GEO |
| **GSE48075** | 142 | | 69 (43-89) | 38 | 33 | 41 | 22 | 8 | 123 | 12 | 119 | 8 | 52 | 17 | OS, DSS | | | [3] | GEO |
| **GSE32894** | 345 | 25 | 71 (20-96) | 116 | 97 | 85 | 7 | 1 | 49 | 22 | | | 228 | 80 | OS | | | [4] | GEO |
| **GSE13507** | 188 | 67 | 66 (24-88) | 24 | 80 | 31 | 19 | 11 | 149 | 15 | 158 | 7 | 135 | 30 | OS | | | [5] | GEO |

**b.** Bulk RNA-seq datasets of pan-tumors

| Datasets | Samples | | | Age (yr) | Tissue | Cancer type | Gender | | Survival | Download |
|---|---|---|---|---|---|---|---|---|---|---|
| | Total | T | N | | | | Male | Female | | |
| **TCGA** | 11123 | 10391 | 732 | 61 (14-90) | 138 | 33 | 5344 | 5739 | OS, PFI, DFI, DSS | TCGA |
| **PCAWG** | 1466 | 1305 | 161 | 61 (17-90) | 22 | 29 | 691 | 654 | OS | UCSC Xena |
| **ICGC** | 8746 | 8097 | 649 | 60 (14-90) | 16 | 21 | 4030 | 4716 | OS | UCSC Xena |
| **TARGET** | 734 | 723 | 11 | 4 (0-30) | | 7 | 321 | 293 | | UCSC Xena |
| **GSE2109** | 2158 | 2158 | | | 192 | | 697 | 1458 | | GEO |

**c.** Baseline bulk RNA-seq datasets of pan-tumors treated with immune-checkpoint blockade therapy

| Datasets | Patients | Cancer type | Treatment | Response | | | OS | PFS | Gender | | Ref. | Download |
|---|---|---|---|---|---|---|---|---|---|---|---|---|
| | | | | R | NR | NE | | | Male | Female | | |
| **IMvigor210** | 348 | BLCA | anti-PDL1 | 68 | 230 | 50 | √ | | 272 | 76 | [6] | R package |
| **Kallisto** | 25 | BLCA | anti-PDL1 | 7 | 14 | 4 | √ | √ | 22 | 3 | [7] | zenodo |
| **GSE111636** | 11 | BLCA | anti-PD1 | 6 | 5 | | | | | | | GEO |

| | | | | | | | | | | | | |
|---|---|---|---|---|---|---|---|---|---|---|---|---|
| **GSE173839** | 71 | BRCA | anti-PDL1+PARPIs | 29 | 42 | | | | | 71 | [8] | GEO |
| **GSE194040** | 69 | BRCA | anti-PD1+Chemotherapy | 31 | 38 | | | | | 69 | [9] | GEO |
| **phs002419** | 14 | BRCA | anti-PD1+Chemotherapy | 4 | 9 | 1 | √ | √ | | 14 | [10] | dbGaP |
| **Checkmate009** | 16 | KIRC | anti-PD1 | 3 | 13 | | √ | √ | 13 | 3 | [11,12] | ArrayExpress |
| **Checkmate010** | 45 | KIRC | anti-PD1 | 11 | 34 | | √ | √ | 30 | 15 | [12,13] | Supplements |
| **Checkmate025** | 120 | KIRC | anti-PD1 | 25 | 86 | 9 | √ | √ | 94 | 26 | [12,14] | EGA |
| **E_MTAB_3218** | 59 | KIRC | anti-PD1 | 13 | 43 | 3 | √ | √ | 39 | 20 | [15] | ArrayExpress |
| **Miao_2018** | 33 | KIRC | anti-PD1/anti-PDL1/anti-PD1+anti-CTLA4 | 8 | 25 | | √ | √ | 24 | 9 | [16] | |
| **GSE67501** | 11 | KIRC | anti-PD1 | 4 | 7 | | | | 7 | 4 | [17] | GEO |
| **IMmotion151** | 407 | KIRC | anti-PDL1+anti-VEGF | 150 | 230 | 27 | | √ | 281 | 126 | [18] | EGA |
| **Javelin101** | 354 | KIRC | anti-PDL1+VEGFRIs | | | | | √ | 257 | 97 | [19] | Supplements |
| **GSE179730** | 11 | COSCC | anti-PD1 | 3 | 8 | | √ | √ | | | [20] | GEO |
| **GSE162137** | 25 | CTCL | anti-PD1 | 11 | 14 | | | | | | [21] | GEO |
| **GSE165252** | 35 | EAC | anti-PD1+Chemoterapy+Radiotherapy | 12 | 20 | 3 | | | | | [22] | GEO |
| **PRJNA482620** | 17 | GBM | anti-PD1 | 10 | 7 | | √ | | | | [23] | NCBI |
| **PRJEB25780** | 45 | GC | anti-PD1 | 12 | 33 | | | | | | [24] | NCBI |
| **GSE195832** | 28 | HNSCC | anti-PD1 | 9 | 19 | | | | 26 | 2 | [25] | Mendeley Data |
| **TJ_Val** | 20 | HNSCC | anti-PD1 | 5 | 15 | | | | | | [25] | Mendeley Data |
| **GSE126044** | 16 | NSCLC | anti-PD1 | 5 | 11 | | | | | | [26] | GEO |
| **GSE135222** | 27 | NSCLC | anti-PD1 | | | | | √ | 22 | 5 | [27] | GEO |
| **OAK** | 344 | NSCLC | anti-PDL1 | 48 | 270 | 26 | √ | √ | 219 | 125 | [28] | EGA |
| **POPLAR** | 95 | NSCLC | anti-PDL1 | 13 | 74 | 8 | √ | √ | 68 | 27 | [28] | EGA |
| **Checkmate038** | 49 | SKCM | anti-PD1 | 9 | 34 | 6 | √ | √ | 24 | 25 | [15] | ArrayExpress |
| **GSE115821** | 14 | SKCM | anti-PD1/ anti-PD1+anti-CTLA4/ anti-CTLA4 | 2 | 12 | | | | | | [29] | GEO |
| **GSE131521** | 17 | SKCM | anti-PD1 | | | | √ | | | | [30] | GEO |
| **GSE78220** | 27 | SKCM | anti-PD1 | 15 | 12 | | √ | | 19 | 8 | [31] | GEO |
| **GSE91061** | 51 | SKCM | anti-PD1 | 10 | 39 | 2 | | | | | [32] | GEO |
| **phs000452** | 116 | SKCM | anti-PD1/ anti-CTLA4 | 48 | 68 | | √ | √ | 67 | 49 | [33] | dbGaP |
| **PRJEB23709** | 73 | SKCM | anti-PD1/ anti-PD1+anti-CTLA4 | 40 | 33 | | √ | √ | 47 | 26 | [34] | NCBI |
| **SRP067586** | 9 | SKCM | anti-CTLA4 | 4 | 5 | | √ | | 5 | 4 | [35] | NCBI |

| | | | | | | | | | |
|---|---|---|---|---|---|---|---|---|---|
| **GSE100797** | 25 | SKCM | IL-2 | 10 | 15 | √ | √ | [36] | GEO |
| **GSE96619** | 5 | SKCM | anti-PD1 | 2 | 3 | | | [37] | GEO |
| **GSE202687** | 9 | UVM | anti-PD1+anti-CTLA4 | | 9 | | | [38] | GEO |

**d.** Single-cell RNA-seq dataset of bladder cancer

| Dataset | Patients Number | Samples Number | Total cells | Tumor cells | CD8Ts | CD4Ts | Monocytes | Macrophages | Treg | Plasma | B cells | Fibroblasts | Endothelial | Ref. | Download |
|---|---|---|---|---|---|---|---|---|---|---|---|---|---|---|---|
| **Salomé's dataset** | 15 | 17 | 70244 | 17884 | 10303 | 8456 | 6383 | 3311 | 8153 | 2037 | 5202 | 3988 | 2034 | [39] | Mendeley Data |

TMB = tumor mutation load

pT/cT = pathological/clinical tumor stage

pN/cN = pathological/clinical lymph node status (0 = no lymph node metastases, 1~3 = lymph node metastases)

pM/cM = pathological/clinical metastasis status (0 = no distant metastases, 1 = distant metastases)

Samples: T = tumor samples, N = benign samples

Survival: OS = overall survival, PFI = progression-free interval, DFI = disease-free interval, DSS = disease-specific survival,
RFS = recurrence-free survival, PFS = progression-free interval

Cancer type: BLCA = bladder cancer, BRCA = breast cancer, KIRC = kidney clear cell carcinoma, COSCC = oral-cavity squamous cell carcinoma,
EAC = esophageal adenocarcinoma, GBM = Glioblastoma, GC = gastric cancer, HNSCC = head and neck squamous cell carcinoma,
NSCLC = non-small cell lung cancer, SKCM = melanoma, UVM = uveal melanoma, CTCL = cutaneous T cell lymphoma

Response: R = response, NR = non-response, NE = not evaluation

**Supplementary Table 2.** Metadata of LY dataset.

| Patient | TIDE Subtype | pT | Pathological Diagnosis | Tumor Volume (mm$^3$) | Ki67 (%) | CK5/6 | PHH3 | CD44 | RNA-seq Samples |
|---|---|---|---|---|---|---|---|---|---|
| **N1** | SII | Ta | High-grade papillary urothelial carcinoma | 3086.25 | 0.70 | - | - | - | 2 |
| **N2** | SII | | Inverted urothelial papilloma | 947.93 | 0.05 | + | - | + | 1 |
| **N3** | SI | Ta | High-grade papillary urothelial carcinoma | | 0.30 | - | - | + | 2 |
| **N4** | SII/SIII | T1 | High-grade invasive urothelial carcinoma | 12539.69 | 0.70 | + | - | - | 2 |
| **N5** | SIII | Tis | Urothelial carcinoma in situ | 300758.62 | 0.70 | - | - | - | 2 |
| **N6** | SI | | Inverted urothelial papilloma | 77.11 | 0.03 | - | - | + | 1 |
| **N7** | SI | Ta | High-grade papillary urothelial carcinoma | 735.00 | 0.30 | - | + | + | 1 |
| **N8** | SI | Ta | High-grade papillary urothelial carcinoma | 487.72 | 0.05 | - | - | + | 1 |
| **N9** | SIII | Ta | High-grade papillary urothelial carcinoma | 6265.83 | | - | - | - | 1 |
| **N10** | SIII | T2 | High-grade papillary urothelial carcinoma | 8485.17 | | + | - | - | 1 |
| **N11** | SII/SIII | T4 | High-grade invasive urothelial carcinoma | 368166.16 | 0.80 | + | - | - | 2 |
| **N12** | SII | T1 | High-grade papillary urothelial carcinoma | 7895.84 | 0.60 | - | - | - | 2 |
| **N13** | SIII | T2 | High-grade invasive urothelial carcinoma | 6845.02 | 0.50 | - | + | - | 2 |
| **N14** | SIII | Tis | Urothelial carcinoma in situ | | 0.60 | + | + | - | 2 |
| **N16** | SI | Ta | Low-grade papillary urothelial carcinoma | 95.51 | 0.05 | - | - | + | 1 |
| **N17** | SII | Ta | Low-grade papillary urothelial carcinoma | 798.33 | 0.10 | - | - | - | 2 |
| **N18** | SII | Ta | Urothelial neoplasms of low malignant potential | | 0.05 | - | - | - | 1 |
| **N19** | SII | T1 | High-grade invasive urothelial carcinoma | 627.49 | 0.30 | - | - | - | 2 |
| **N20** | SII | Ta | High-grade papillary urothelial carcinoma | | 0.50 | + | - | - | 1 |
| **N21** | SI | Ta | Low-grade papillary urothelial carcinoma | 402.65 | 0.30 | - | - | - | 2 |

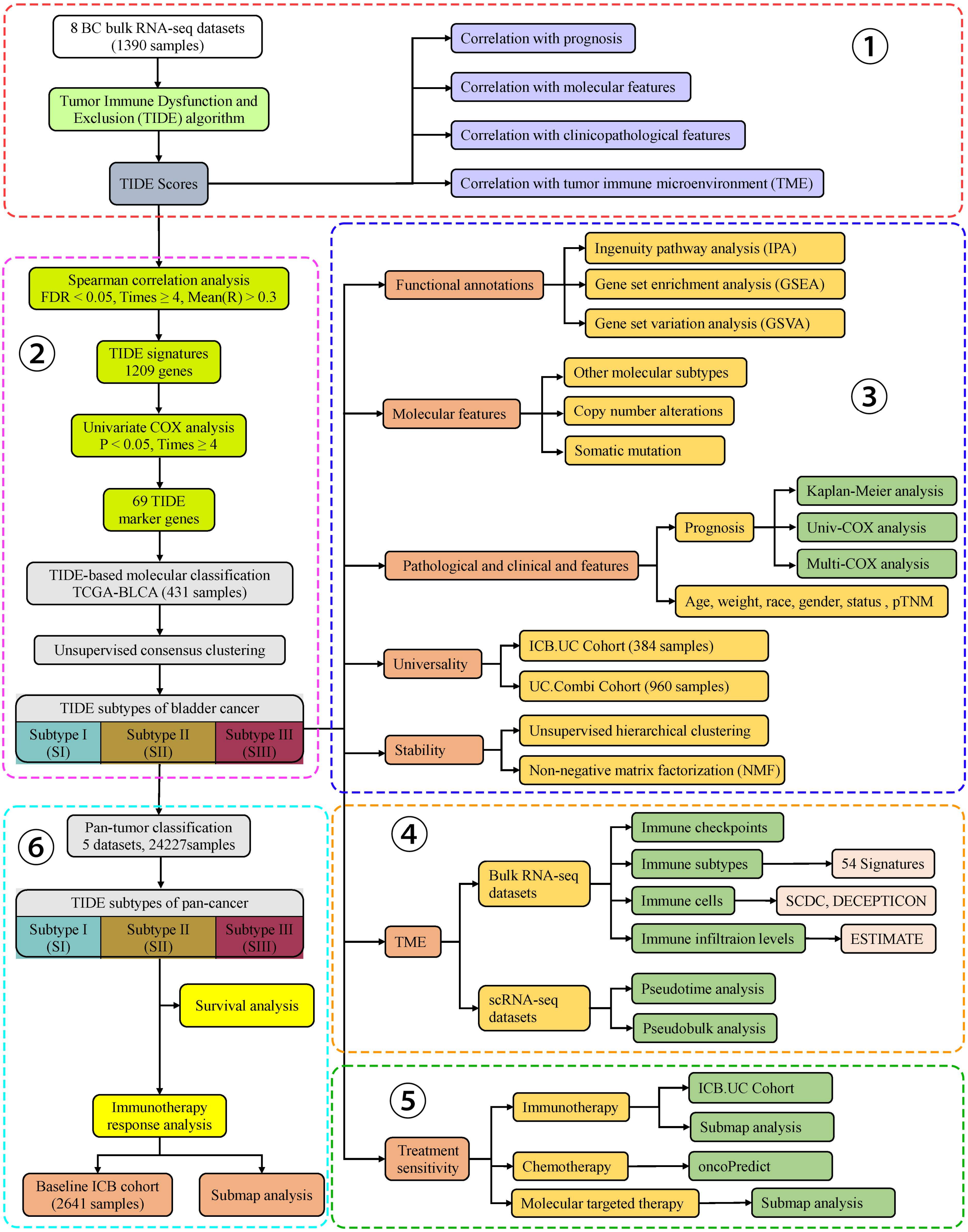


**Supplementary Figure 1. The overall design of the current study.**

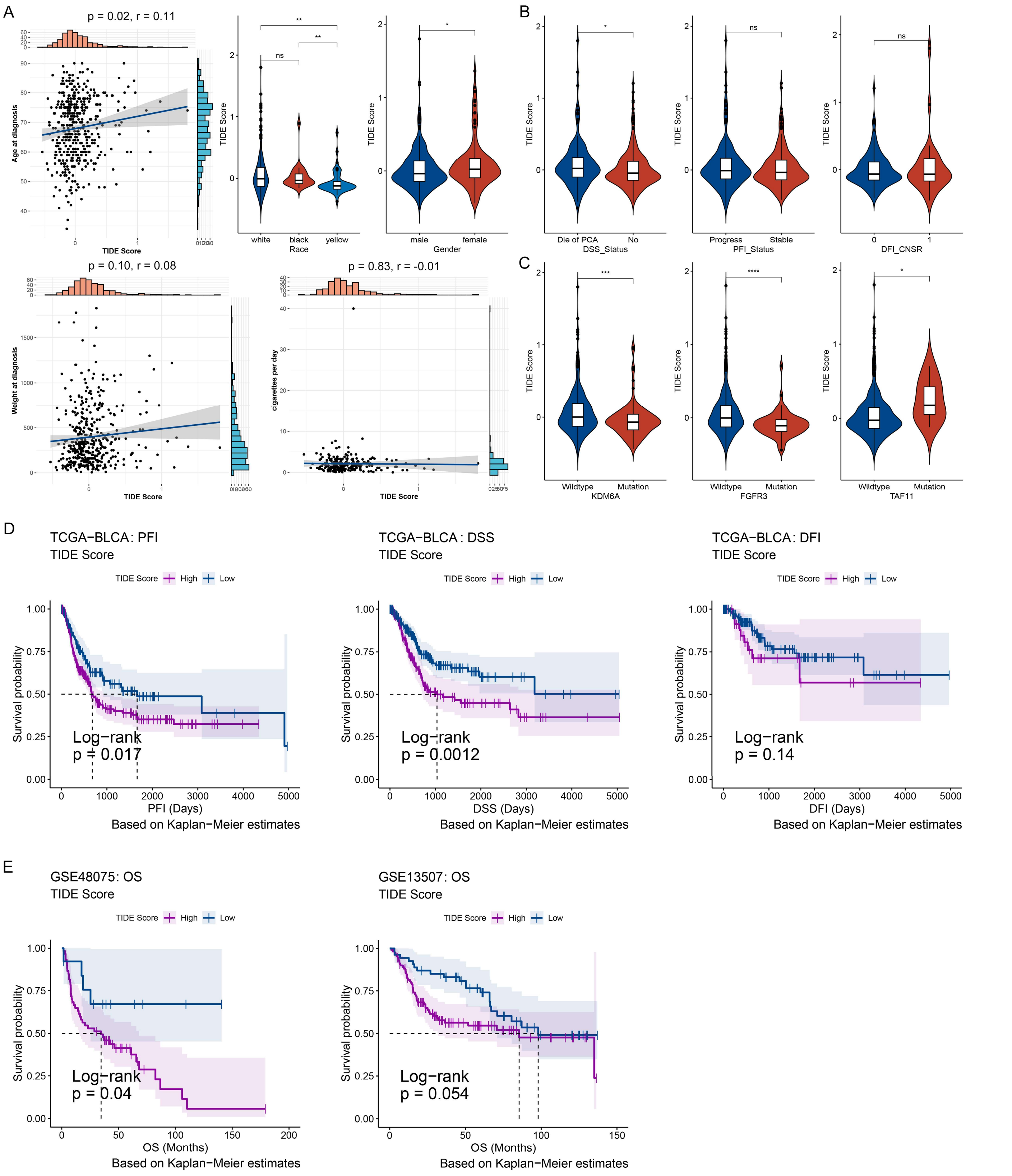


**Supplementary Figure 2. Correlations of TIDE status with clinicopathological and molecular features in the BC patients** (related to Figure 1). **(A)** Correlations of TIDE scores with age and weight at diagnosis, race, gender, and daily smoking in BC patients. **(B)** Comparisons of TIDE scores with disease specific death (DSS_Status), progress (PFI_Status) and disease presence (DFI_Status). **(C)** Comparisons of TIDE scores between mutant and wild-type patients at KDM6A, FGFR3 and TAF11 genes. **(D, E)** Kaplan-Meier (K-M) analysis demonstrated a correlation between TIDE scores and the prognosis of BC patients from TCGA-BLCA **(D)**, GSE48075 and GSE13507 **(E)** datasets. OS, overall survival; PFI, progression-free interval; DFI, disease-free interval; DSS, disease-specific survival. Dashed line: median survival time. Color range: 95% confidence interval (CI). *$p < 0.05$, **$p < 0.01$, ***$p < 0.001$, ****$p < 0.0001$; ns, no significance.

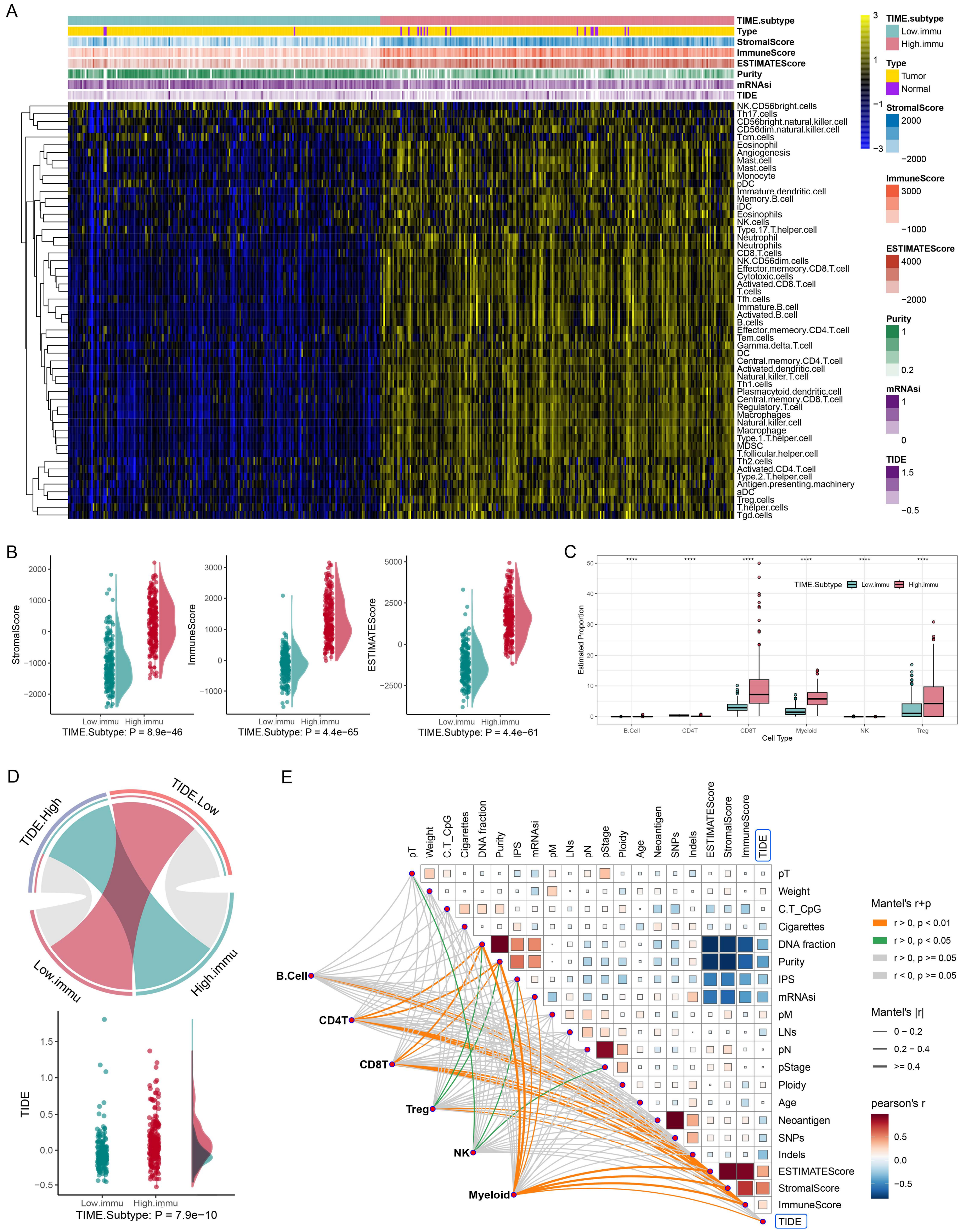

**Supplementary Figure 3. Correlations of TIDE status with TIME in the BC patients. (A)** Consensus clustering divided BC patients from TCGA-BLCA into two subtypes according to the activity scores of 54 published immune signatures: low immune infiltration (Low.immu) and high immune infiltration (High.immu) subtypes. **(B)** Raincloudplot shows the TME scores (StromalScore, ImmuneScore, ESTIMATEScore) of BC patients in two TIME subtypes. **(C)** Comparisons of immunocyte abundance among the TIME subtypes of TCGA-BLCA. Cell proportions are assessed by the DECEPTICON algorithm. **(D)** The association between TIME subtypes and TIDE groups, achieved by the hypergeometric test. Gray lines represent no significance. The TIDE groups were dichotomized at the median TIDE scores **(Upper)**. Raincloud plot showing the TIDE levels in BC patients among the TIME subtypes **(Under)**. **(E)** Heatmaps showing correlations of TIDE scores with clinical and molecular features, and tumor microenvironment (TME) scores (Stromal, Immune, ESTIMATE scores), achieved by Pearson correlation analysis. Links showing the correlations between immunocyte abundance and TIDE scores, TME scores, and clinical and molecular features, implemented by Mantel test. IPS, Immunophenotype score; LNs, Number of lymph nodes examined; SNPs, single nucleotide polymorphisms. The square size represents the absolute value of Pearson's R.

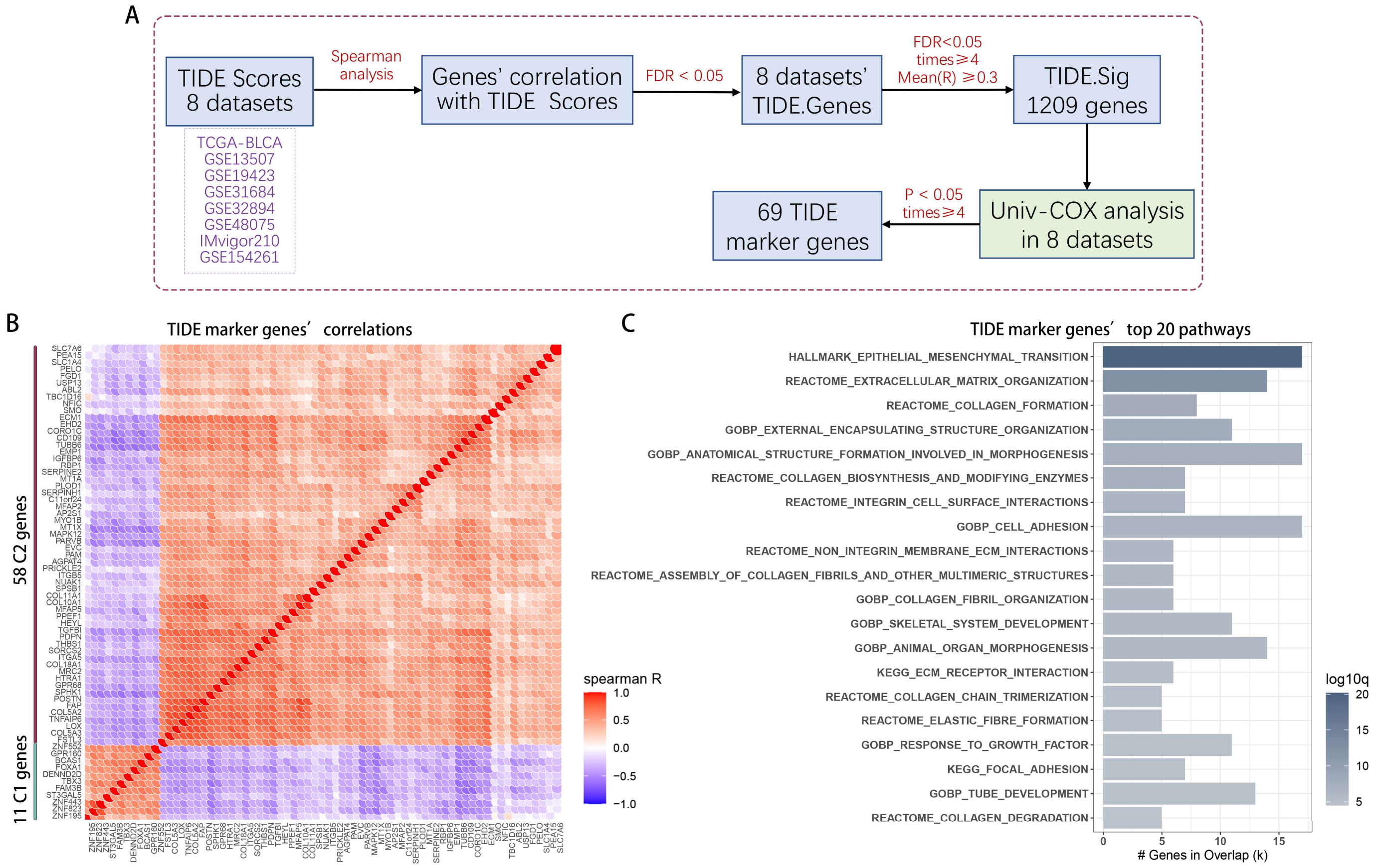


**Supplementary Figure 4. Identification of TIDE marker genes for molecular subtyping. (A)** The workflow of TIDE marker genes identification. **(B)** Correlation heatmap of the 69 TIDE marker genes. These genes are mainly divided into two clusters: C1 consists of 11 genes and C2 comprises 58 genes. **(C)** Top 20 signaling pathways enriched based on the 58 C2 genes.

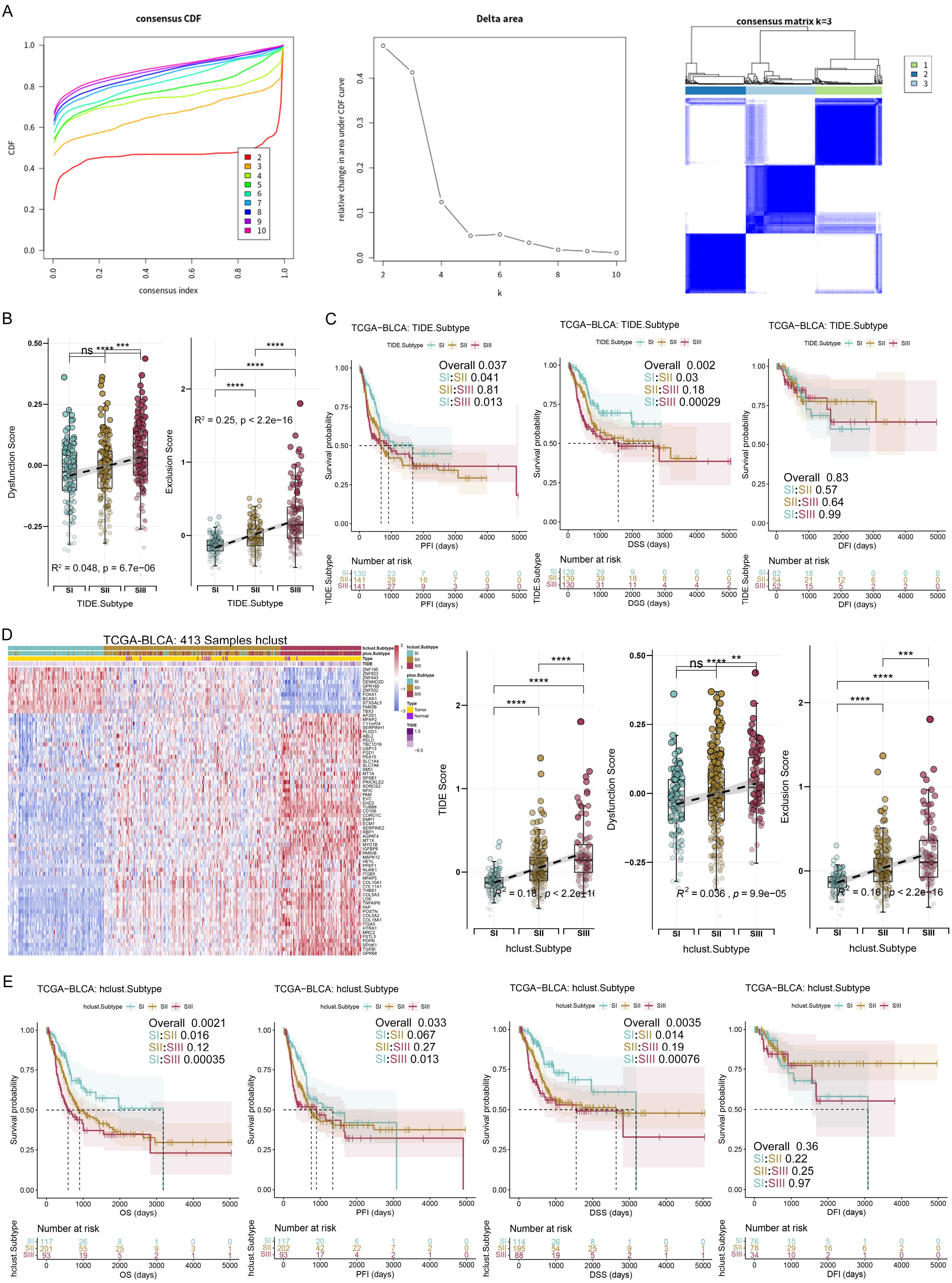
A
consensus CDF
CDF
consensus index
Delta area
relative change in area under CDF curve
k
consensus matrix k=3
B
Dysfunction Score
TIDE.Subtype
R² = 0.048, p = 6.7e−06
Exclusion Score
R² = 0.25, p < 2.2e−16
C
TCGA−BLCA: TIDE.Subtype
Survival probability
PFI (days)
Overall 0.037
SI:SII 0.041
SII:SIII 0.81
SI:SIII 0.013
DSS (days)
Overall 0.002
SI:SII 0.03
SII:SIII 0.18
SI:SIII 0.00029
DFI (days)
Overall 0.83
SI:SII 0.57
SII:SIII 0.64
SI:SIII 0.99
Number at risk
D
TCGA-BLCA: 413 Samples hclust
TIDE Score
R² = 0.18, p < 2.2e−16
Dysfunction Score
R² = 0.036, p = 9.9e−05
Exclusion Score
R² = 0.16, p < 2.2e−16
hclust.Subtype
E
TCGA−BLCA: hclust.Subtype
OS (days)
Overall 0.0021
SI:SII 0.016
SII:SIII 0.12
SI:SIII 0.00035
PFI (days)
Overall 0.033
SI:SII 0.067
SII:SIII 0.27
SI:SIII 0.013
DSS (days)
Overall 0.0035
SI:SII 0.014
SII:SIII 0.19
SI:SIII 0.00076
DFI (days)
Overall 0.36
SI:SII 0.22
SII:SIII 0.25
SI:SIII 0.97
Number at risk

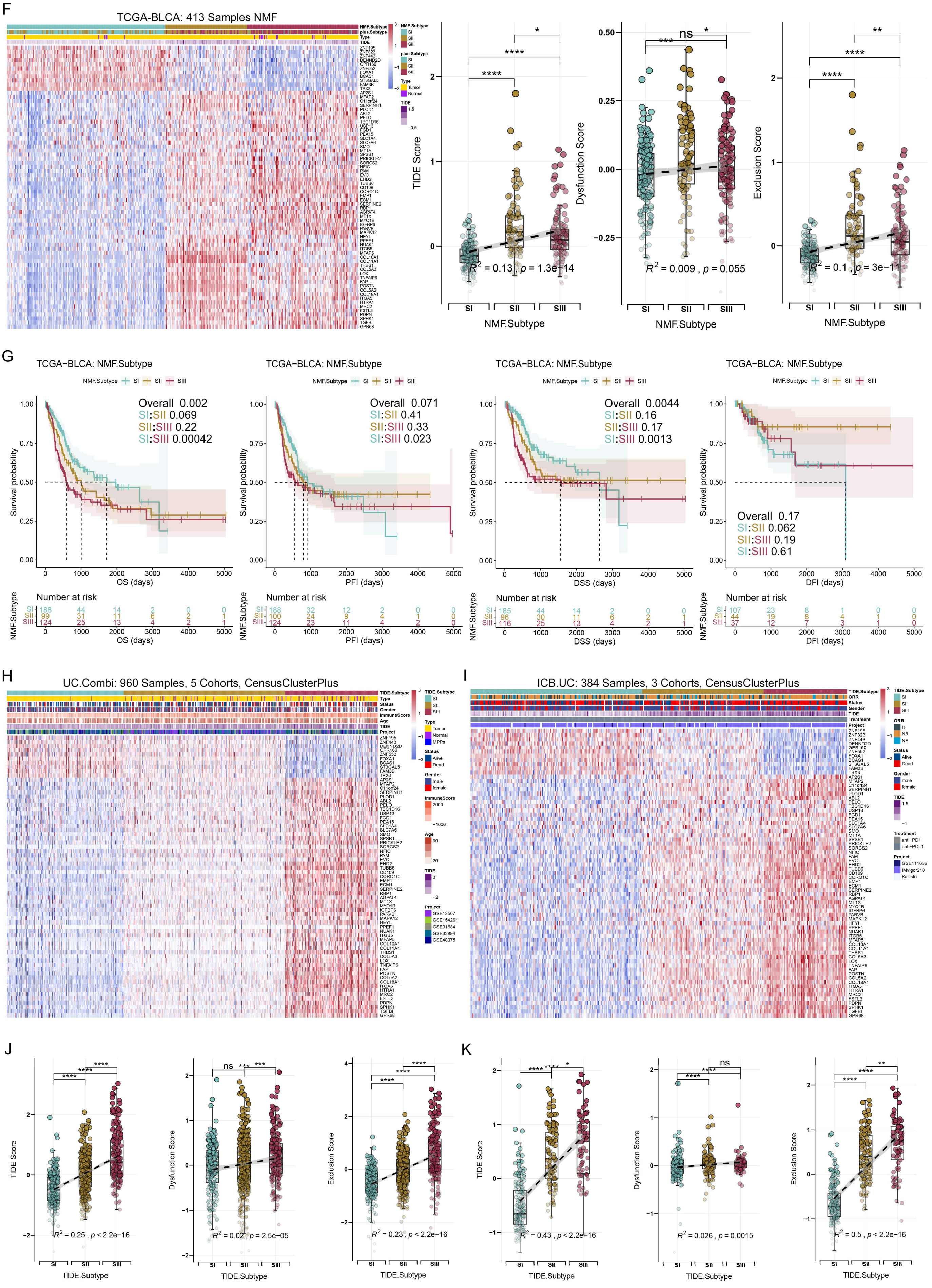
F
TCGA-BLCA: 413 Samples NMF
NMF.Subtype
TIDE Score
Dysfunction Score
Exclusion Score
G
TCGA−BLCA: NMF.Subtype
Overall 0.002
SI:SII 0.069
SII:SIII 0.22
SI:SIII 0.00042
Overall 0.071
SI:SII 0.41
SII:SIII 0.33
SI:SIII 0.023
Overall 0.0044
SI:SII 0.16
SII:SIII 0.17
SI:SIII 0.0013
Overall 0.17
SI:SII 0.062
SII:SIII 0.19
SI:SIII 0.61
Survival probability
OS (days)
PFI (days)
DSS (days)
DFI (days)
Number at risk
H
UC.Combi: 960 Samples, 5 Cohorts, CensusClusterPlus
I
ICB.UC: 384 Samples, 3 Cohorts, CensusClusterPlus
J
K
TIDE.Subtype

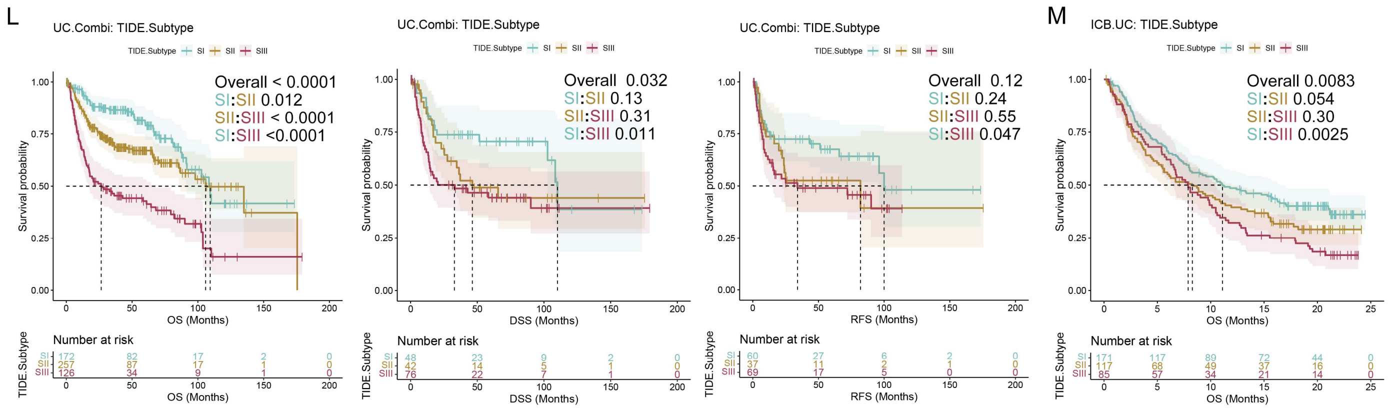

**Supplementary Figure 5. Identifications of three BC TIDE subtypes based on TIDE marker genes** (related to Figure 2). **(A)** Cumulative distribution function (CDF) curves of the consensus score from k = 2 to 10 (left). The relative change in the area under the CDF curve from k = 2 to 10 (**medium**). Consensus matrix for k = 3, which was the optimal cluster number (**right**). **(B)** Levels and trends of Dysfunction scores and Exclusion scores among the TIDE subtypes. **(C)** K-M analysis shows significant differences in PFI and DSS among the TIDE subtypes of TCGA-BLCA. **(D)** Unsupervised hierarchical clustering (hclust) based on the expression of the 69 TIDE marker genes classified TCGA-BLCA patients into three subtypes: Subtype I (SI), Subtype II (SII), and Subtype III (SIII) **(left)**. Levels and trends of TIDE, Dysfunction and Exclusion scores among the TIDE subtypes **(right)**. **(E)** K-M analysis of the TIDE subtypes of TCGA-BLCA classified by hclust. **(F)** Unsupervised nonnegative matrix factorization (NMF) based on the expression of the 69 TIDE marker genes classified TCGA-BLCA patients into three subtypes: SI, SII and SIII **(left)**. Levels and trends of TIDE, Dysfunction and Exclusion scores among the TIDE subtypes **(right)**. **(G)** K-M analysis of the TIDE subtypes of TCGA-BLCA classified by NMF. **(H, I)** Consensus clustering based on the expression of 69 TIDE marker genes classified UC.Combi **(H)** and ICB.UC **(I)** cohorts into three subtypes. **(J, K)** Levels and trends of TIDE, Dysfunction and Exclusion scores among the TIDE subtypes of UC.Combi **(J)** and ICB.UC **(K)** cohorts. **(L, M)** K-M analysis of the TIDE subtypes of UC.Combi **(L)** and ICB.UC **(M)** cohorts. *p < 0.05, **p < 0.01, ***p < 0.001, ****p < 0.0001; ns, no significance.

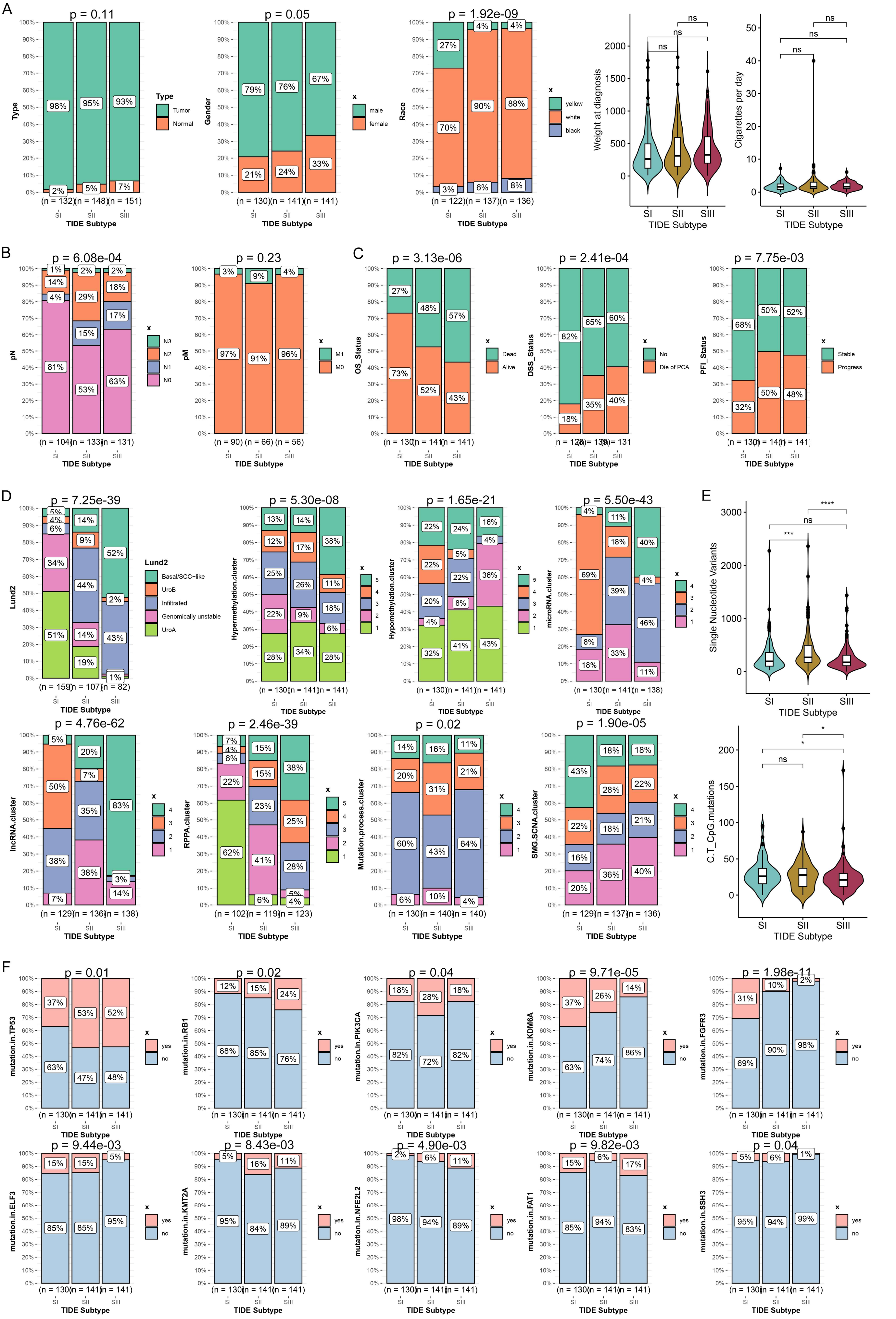
A
p = 0.11
Type
Tumor
Normal
98%
95%
93%
2%
5%
7%
(n = 132)
(n = 148)
(n = 151)
TIDE Subtype
p = 0.05
Gender
male
female
79%
76%
67%
21%
24%
33%
(n = 130)
(n = 141)
(n = 141)
p = 1.92e-09
Race
yellow
white
black
27%
4%
4%
70%
90%
88%
3%
6%
8%
(n = 122)
(n = 137)
(n = 136)
Weight at diagnosis
Cigarettes per day
ns
SI
SII
SIII
B
p = 6.08e-04
pN
N3
N2
N1
N0
1%
2%
2%
14%
29%
18%
4%
17%
15%
81%
53%
63%
(n = 104)
(n = 133)
(n = 131)
p = 0.23
pM
M1
M0
3%
9%
4%
97%
91%
96%
(n = 90)
(n = 66)
(n = 56)
C
p = 3.13e-06
OS_Status
Dead
Alive
27%
48%
57%
73%
52%
43%
(n = 130)
(n = 141)
(n = 141)
p = 2.41e-04
DSS_Status
No
Die of PCA
82%
65%
60%
18%
35%
40%
p = 7.75e-03
PFI_Status
Stable
Progress
68%
50%
52%
32%
50%
48%
D
p = 7.25e-39
Lund2
Basal/SCC−like
UroB
Infiltrated
Genomically unstable
UroA
5%
4%
6%
14%
9%
52%
34%
44%
2%
51%
14%
43%
19%
1%
(n = 159)
(n = 107)
(n = 82)
p = 5.30e-08
Hypermethylation.cluster
13%
14%
12%
17%
38%
25%
26%
11%
22%
9%
18%
6%
28%
34%
28%
(n = 130)
(n = 141)
(n = 141)
p = 1.65e-21
Hypomethylation.cluster
22%
24%
16%
4%
22%
5%
36%
22%
20%
8%
4%
41%
43%
32%
p = 5.50e-43
microRNA.cluster
4%
11%
18%
40%
69%
4%
39%
46%
8%
33%
18%
11%
(n = 130)
(n = 141)
(n = 138)
E
Single Nucleotide Variants
****
ns
***
p = 4.76e-62
lncRNA.cluster
5%
20%
50%
7%
35%
83%
38%
38%
3%
14%
7%
(n = 129)
(n = 136)
(n = 138)
p = 2.46e-39
RPPA.cluster
7%
4%
6%
15%
38%
22%
15%
23%
25%
62%
41%
28%
6%
5%
4%
(n = 102)
(n = 119)
(n = 123)
p = 0.02
Mutation.process.cluster
14%
16%
11%
20%
31%
21%
60%
43%
64%
6%
10%
4%
(n = 130)
(n = 140)
(n = 140)
p = 1.90e-05
SMG.SCNA.cluster
43%
18%
18%
28%
22%
22%
18%
21%
16%
36%
40%
20%
(n = 129)
(n = 137)
(n = 136)
C.T_CpG.mutations
*
F
p = 0.01
mutation.in.TP53
yes
no
37%
53%
52%
63%
47%
48%
p = 0.02
mutation.in.RB1
12%
15%
24%
88%
85%
76%
p = 0.04
mutation.in.PIK3CA
18%
28%
18%
82%
72%
82%
p = 9.71e-05
mutation.in.KDM6A
37%
26%
14%
63%
74%
86%
p = 1.98e-11
mutation.in.FGFR3
31%
10%
2%
69%
90%
98%
p = 9.44e-03
mutation.in.ELF3
15%
15%
5%
85%
85%
95%
p = 8.43e-03
mutation.in.KMT2A
5%
16%
11%
95%
84%
89%
p = 4.90e-03
mutation.in.NFE2L2
2%
6%
11%
98%
94%
89%
p = 9.82e-03
mutation.in.FAT1
15%
6%
17%
85%
94%
83%
p = 0.04
mutation.in.SSH3
5%
6%
1%
95%
94%
99%
(n = 130)
(n = 141)
(n = 141)
TIDE Subtype

**Supplementary Figure 6. Comparisons of clinicopathological and molecular features among the TIDE subtypes of BC** (related to Figure 3). (A) Comparisons of sample type, gender, race, weight at diagnosis, and daily smoking among three TIDE subtypes. (B) The proportions of pathological N stage (pN) and pathological M stage (pM) among the TIDE subtypes. (C) The proportions of OS_Status, DSS_Status and DFI_Status among the TIDE subtypes. (D) Associations and similarities of TIDE subtypes with other molecular subtypes. (E) Comparisons of single nucleotide variants **(upper)** and C/T CpG mutations **(under)** among the TIDE subtypes. (F) Comparisons of mutations in commonly identified biomarkers among the three TIDE subtypes in BC patients. *p < 0.05, ***p < 0.001, ****p < 0.0001; ns, no significance.

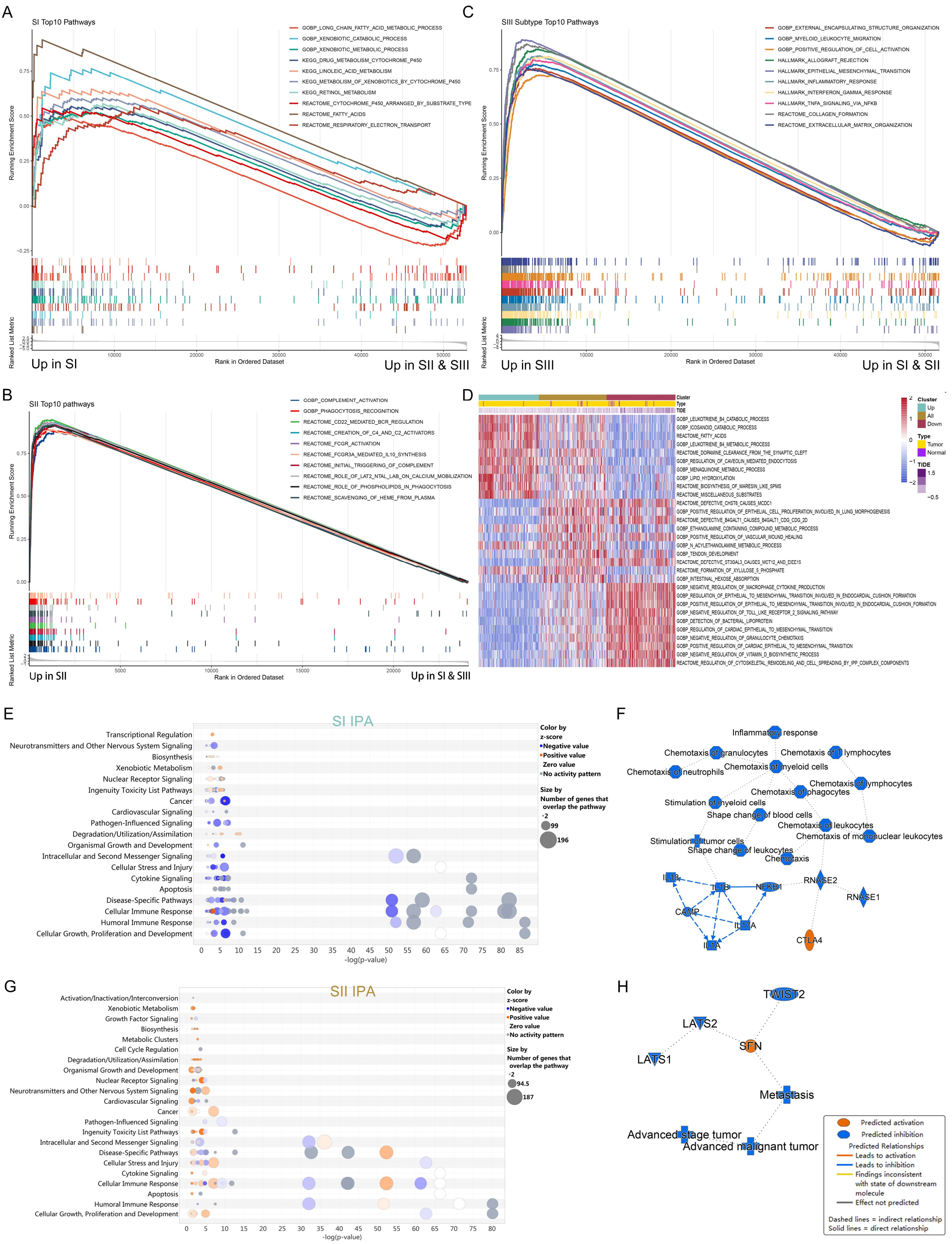

A
SI Top10 Pathways
GOBP_LONG_CHAIN_FATTY_ACID_METABOLIC_PROCESS
GOBP_XENOBIOTIC_CATABOLIC_PROCESS
GOBP_XENOBIOTIC_METABOLIC_PROCESS
KEGG_DRUG_METABOLISM_CYTOCHROME_P450
KEGG_LINOLEIC_ACID_METABOLISM
KEGG_METABOLISM_OF_XENOBIOTICS_BY_CYTOCHROME_P450
KEGG_RETINOL_METABOLISM
REACTOME_CYTOCHROME_P450_ARRANGED_BY_SUBSTRATE_TYPE
REACTOME_FATTY_ACIDS
REACTOME_RESPIRATORY_ELECTRON_TRANSPORT
Running Enrichment Score
Ranked List Metric
Rank in Ordered Dataset
Up in SI
Up in SII & SIII
C
SIII Subtype Top10 Pathways
GOBP_EXTERNAL_ENCAPSULATING_STRUCTURE_ORGANIZATION
GOBP_MYELOID_LEUKOCYTE_MIGRATION
GOBP_POSITIVE_REGULATION_OF_CELL_ACTIVATION
HALLMARK_ALLOGRAFT_REJECTION
HALLMARK_EPITHELIAL_MESENCHYMAL_TRANSITION
HALLMARK_INFLAMMATORY_RESPONSE
HALLMARK_INTERFERON_GAMMA_RESPONSE
HALLMARK_TNFA_SIGNALING_VIA_NFKB
REACTOME_COLLAGEN_FORMATION
REACTOME_EXTRACELLULAR_MATRIX_ORGANIZATION
Up in SIII
Up in SI & SII
B
SII Top10 pathways
GOBP_COMPLEMENT_ACTIVATION
GOBP_PHAGOCYTOSIS_RECOGNITION
REACTOME_CD22_MEDIATED_BCR_REGULATION
REACTOME_CREATION_OF_C4_AND_C2_ACTIVATORS
REACTOME_FCGR_ACTIVATION
REACTOME_FCGR3A_MEDIATED_IL10_SYNTHESIS
REACTOME_INITIAL_TRIGGERING_OF_COMPLEMENT
REACTOME_ROLE_OF_LAT2_NTAL_LAB_ON_CALCIUM_MOBILIZATION
REACTOME_ROLE_OF_PHOSPHOLIPIDS_IN_PHAGOCYTOSIS
REACTOME_SCAVENGING_OF_HEME_FROM_PLASMA
Up in SII
Up in SI & SIII
D
Cluster
Type
TIDE
Up
All
Down
Tumor
Normal
E
SI IPA
Color by z-score
Negative value
Positive value
Zero value
No activity pattern
Size by Number of genes that overlap the pathway
-log(p-value)
F
Inflammatory response
Chemotaxis of granulocytes
Chemotaxis of T lymphocytes
Chemotaxis of neutrophils
Chemotaxis of myeloid cells
Chemotaxis of lymphocytes
Chemotaxis of phagocytes
Stimulation of myeloid cells
Shape change of blood cells
Chemotaxis of leukocytes
Chemotaxis of mononuclear leukocytes
Stimulation of tumor cells
Shape change of leukocytes
Chemotaxis
IL18
IL1B
NFKB1
RNASE2
RNASE1
CAMP
IL17A
IL1A
CTLA4
G
SII IPA
H
TWIST2
LATS2
LATS1
SFN
Metastasis
Advanced stage tumor
Advanced malignant tumor
Predicted activation
Predicted inhibition
Predicted Relationships
Leads to activation
Leads to inhibition
Findings inconsistent with state of downstream molecule
Effect not predicted
Dashed lines = indirect relationship
Solid lines = direct relationship

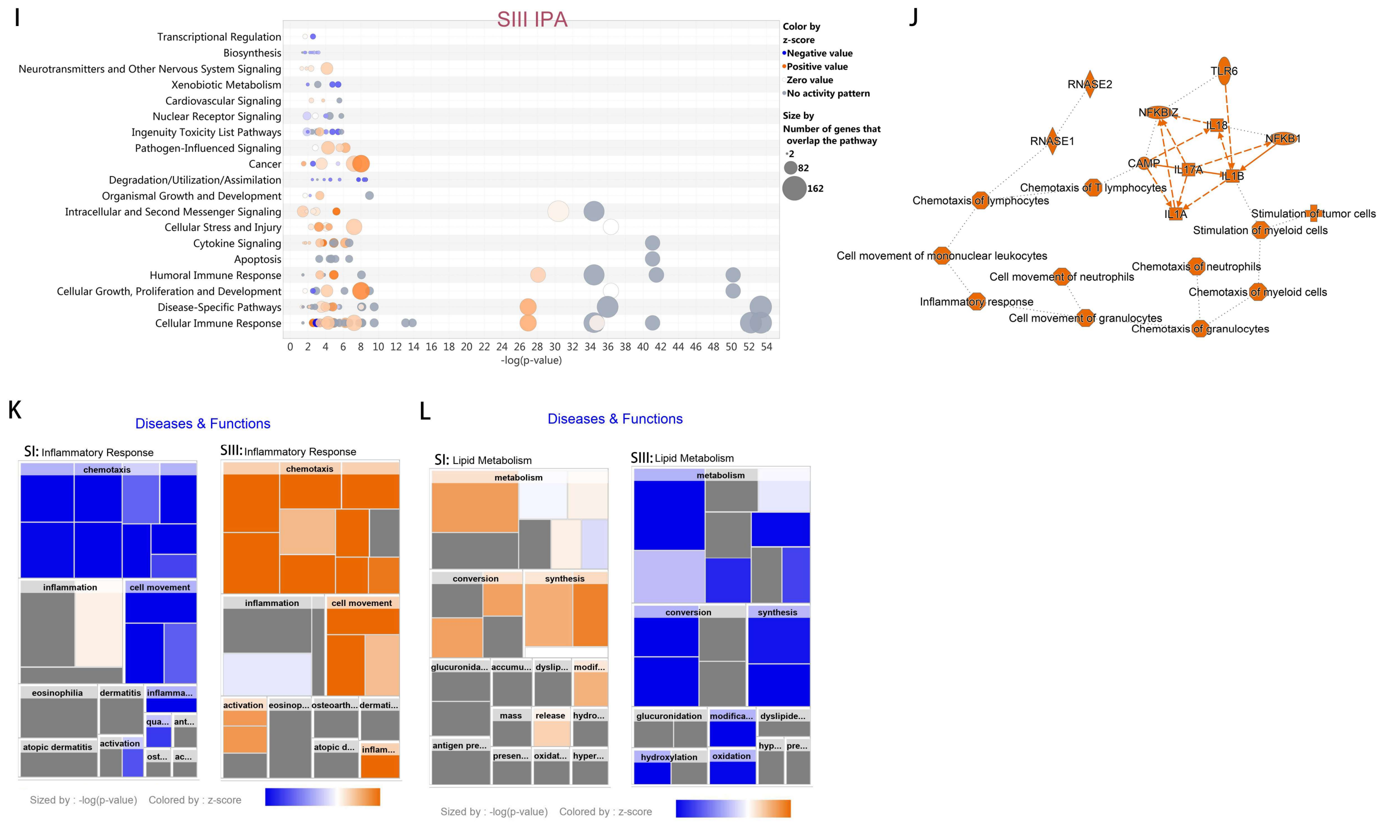


**Supplementary Figure 7. Signaling pathways and functional annotations of three TIDE subtypes of BC**. **(A-C)** Gene set enrichment analysis (GSEA) shows the top 10 signaling pathways enriched in SI **(A)**, SII **(B)** and SIII **(C)**. **(D)** Gene set variation analysis (GSVA) reveals the top 10 signaling pathways enriched in the three TIDE subtypes. **(E, G, I)** Ingenuity pathways analysis (IPA) demonstrates the activation or inhibition status of canonical signaling pathways for SI **(E)**, SII **(G)** and SIII **(I)**. **(F, H, J)** Graphical summary explains biomolecular interactions and biological processes in SI **(F)**, SII **(H)** and SIII **(J)**. **(K, L)** Heatmap showing correlations of marker genes of three TIDE subtype with inflammatory response **(K)** and lipid metabolism **(L)**.

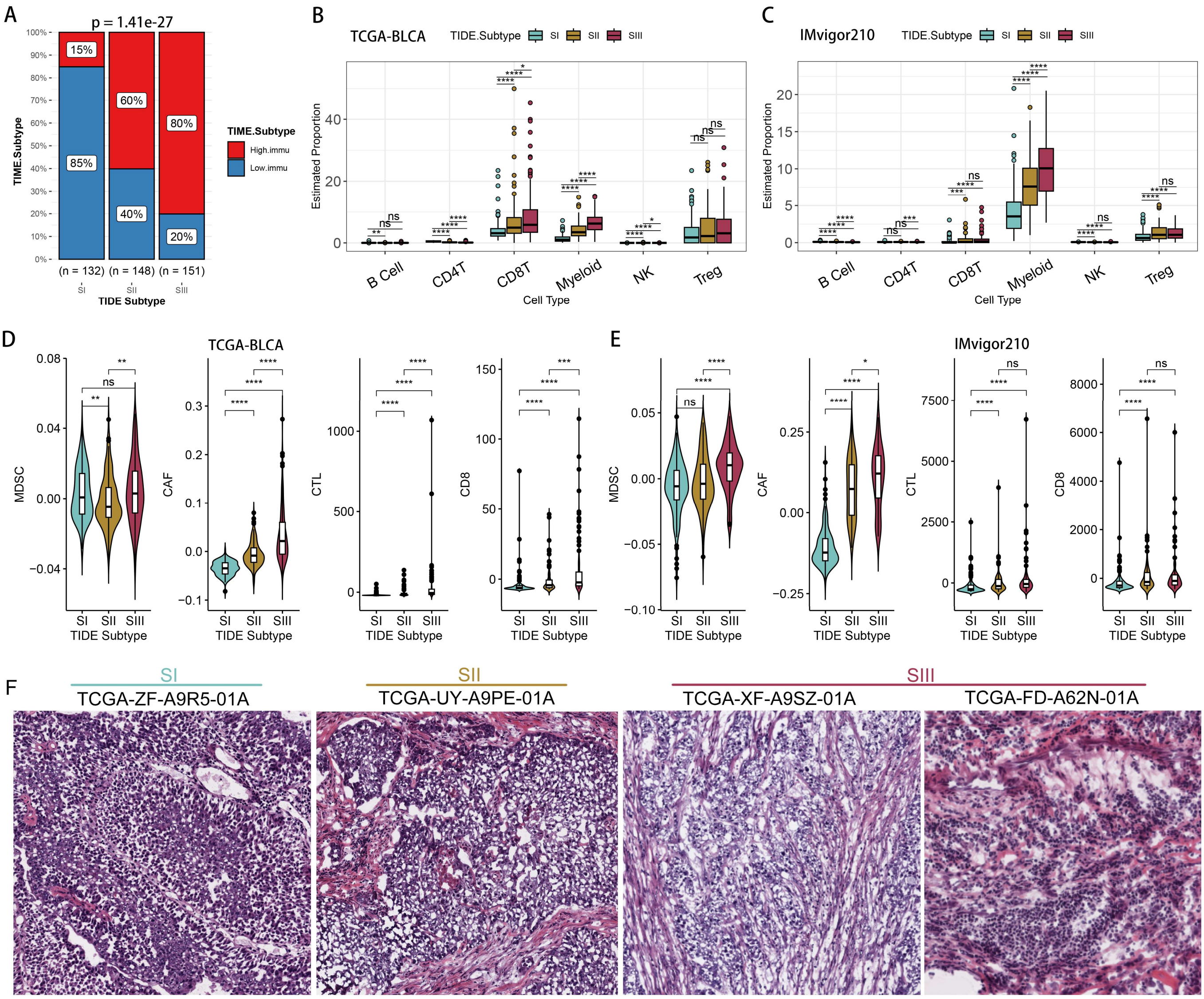


**Supplementary Figure 8. Characterizations of TME patterns among the TIDE subtypes based on bulk RNA-seq datasets** (related to Figure 5). **(A)** Stacked histogram showing the compositional differences of the TIME subtypes between the three TIDE subtypes of BC. **(B, C)** Comparisons of immunocyte abundance among the three TIDE subtypes of TCGA-BLCA **(B)** and IMvigor210 **(C)**. Cell proportions are assessed by the DECEPTICON algorithm. **(D, E)** Comparison of the immunocyte proportion among three TIDE subtypes of TCGA-BLCA **(D)** and IMvigor210 **(E)**. Cell proportions are assessed by the TIDE algorithm. **(F)** Representative TCGA-BLCA H&E histological images of the three subtypes. *p < 0.05, **p < 0.01, ***p < 0.001, ****p < 0.0001; ns, no significance.

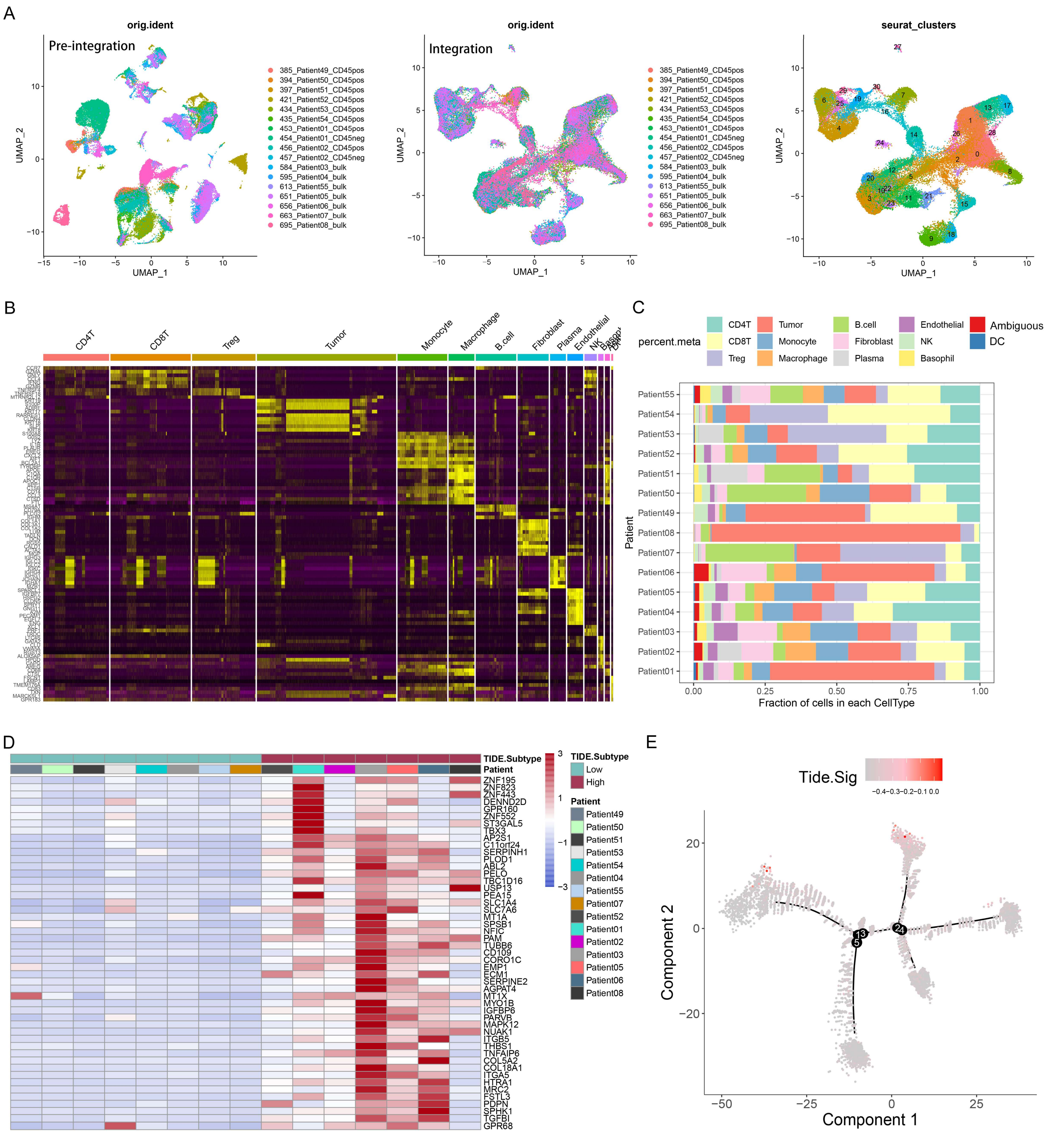


**Supplementary Figure 9. Characterizations of TME patterns among the TIDE subtypes based on single-cell RNA-seq dataset** (related to Figure 6). **(A)** Uniform Manifold Approximation and Projection (UMAP) plot was used to analyze the single-cell RNA-seq dataset. Each color coded for the 17 pre-integrated samples **(left)**, 17 integrated samples **(middle)**, and 31 clusters **(right)** in Salomé's dataset. **(B)** Heatmap of top 10 marker gene expression of the 13 cell types. **(C)** Proportions of the 13 cell types in each sample. **(D)** Consensus clustering based on the expression of 69 TIDE marker genes classified pseudobulk data of Salomé's dataset into two subtypes: Low and High subtypes. **(E)** Expression levels of TIDE marker genes during the CD8T+ T cells (CD8Ts) differentiation trajectory in BC.

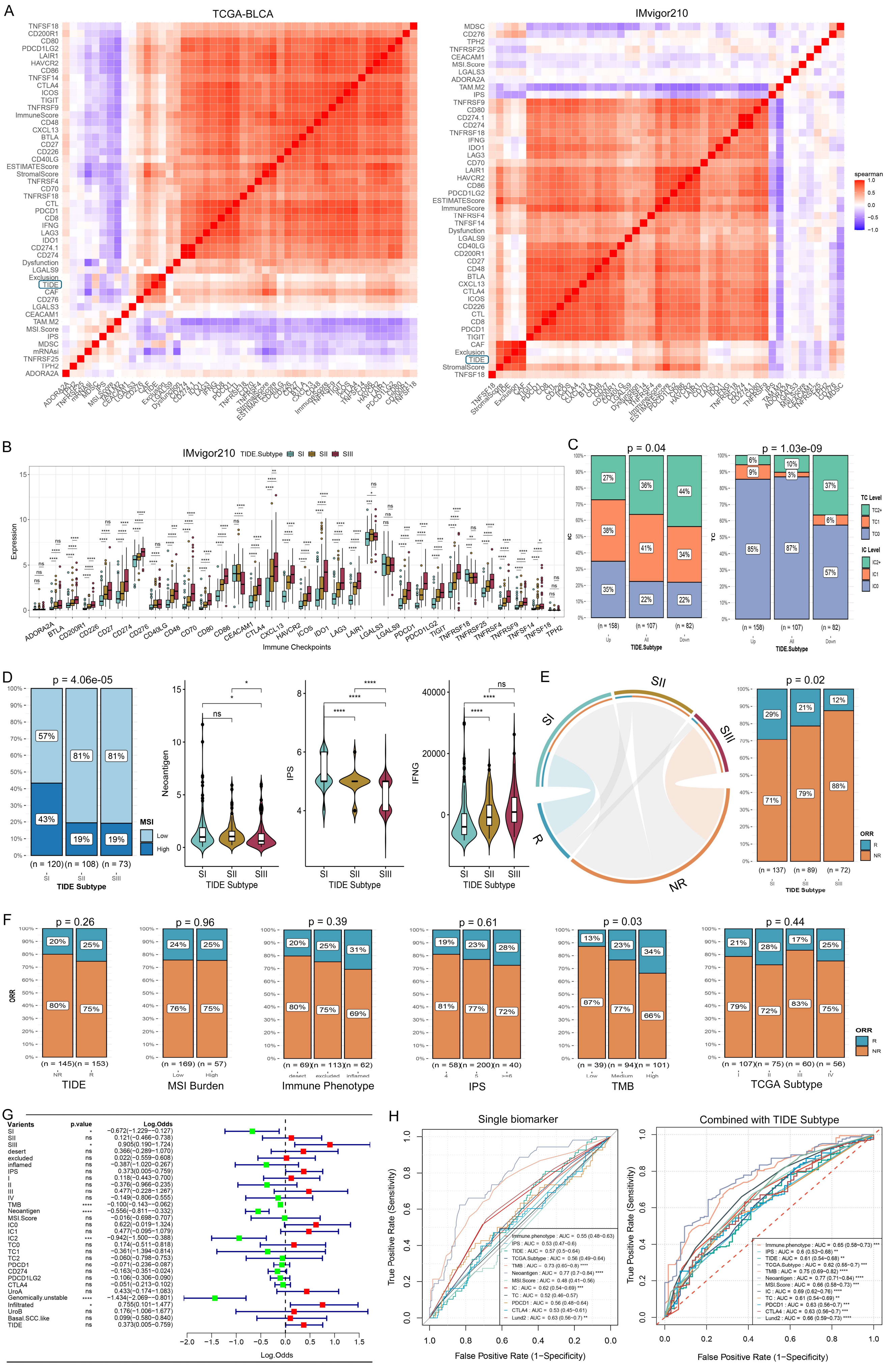
A
TCGA-BLCA
IMvigor210
spearman
B
IMvigor210
TIDE.Subtype SI SII SIII
Expression
Immune Checkpoints
C
p = 0.04
p = 1.03e-09
TC Level
IC Level
TIDE.Subtype
D
p = 4.06e-05
MSI
Low
High
TIDE Subtype
Neoantigen
IPS
IFNG
E
p = 0.02
ORR
R
NR
F
p = 0.26
p = 0.96
p = 0.39
p = 0.61
p = 0.03
p = 0.44
TIDE
MSI Burden
Immune Phenotype
IPS
TMB
TCGA Subtype
G
Varients
p.value
Log.Odds
H
Single biomarker
Combined with TIDE Subtype
True Positive Rate (Sensitivity)
False Positive Rate (1−Specificity)

**Supplementary Figure 10. TIDE subtypes were closely related to ICB response** (related to Figure 7). **(A)** Heatmap showing the correlations of the TIDE scores with expression levels of immune checkpoint molecules from TCGA-BLCA **(left)** and IMvigor210 **(right)** datasets. **(B)** Comparisons of the immune checkpoint molecules expression levels among the TIDE subtypes of IMvigor210 dataset. **(C)** Proportions of IC and TC levels in the TIDE subtypes. **(D)** Comparisons of MSI, neoantigen load, IPS and interferon gamma among three TIDE subtypes of IMvigor210 dataset. **(E)** Hypergeometric test revealed an association between TIDE subtypes of IMvigor210 and ICB responses **(left)**, gray lines represent no significance; Stacked histogram showing the differences of ICB responses among the TIDE subtypes of IMvigor210 dataset **(right)**. **(F)** The relationship of common biomarkers with ICB responses in the IMvigor210 dataset. **(G)** Impacts of the TIDE subtypes and the predictive biomarkers on ICB efficacy, which was achieved by univariate logistic regression analysis. **(H)** ROC curves of single biomarkers (left) and TIDE subtypes+ other biomarkers (right) for predicting the ICB efficacy. R, response; NR, non-response. *$p < 0.05$, **$p < 0.01$, ***$p < 0.001$, ****$p < 0.0001$; ns, no significance.

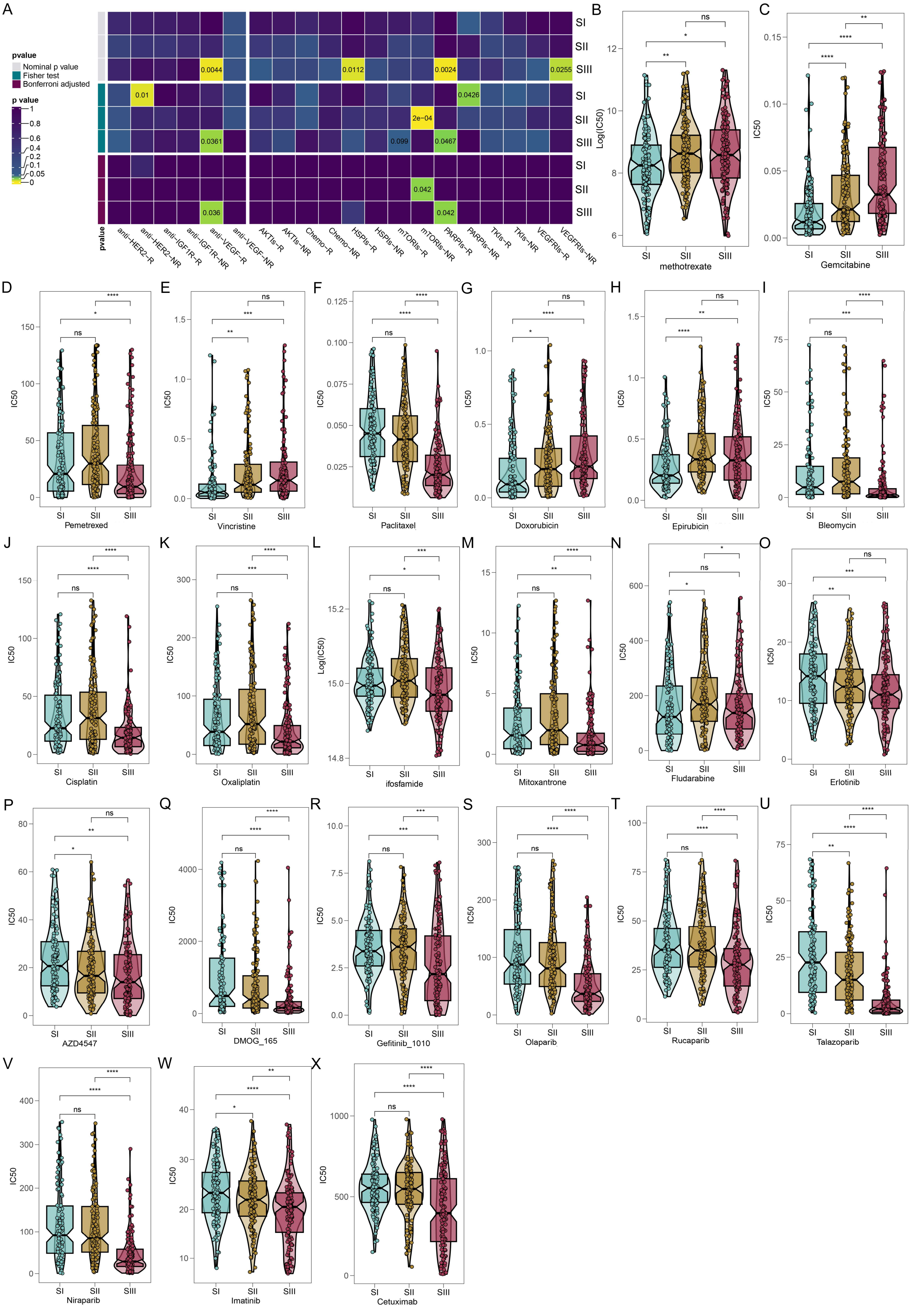
A
pvalue
Nominal p value
Fisher test
Bonferroni adjusted
p value
1
0.8
0.6
0.4
0.2
0.1
0.05
0
0.0044
0.0112
0.0024
0.0255
0.01
0.0426
2e−04
0.0361
0.099
0.0467
0.042
0.036
0.042
SI
SII
SIII
pvalue
anti−HER2−R
anti−HER2−NR
anti−IGF1R−R
anti−IGF1R−NR
anti−VEGF−R
anti−VEGF−NR
AKTIs−R
AKTIs−NR
Chemo−R
Chemo−NR
HSPIs−R
HSPIs−NR
mTORIs−R
mTORIs−NR
PARPIs−R
PARPIs−NR
TKIs−R
TKIs−NR
VEGFRIs−R
VEGFRIs−NR
B
Log(IC50)
methotrexate
C
IC50
Gemcitabine
D
Pemetrexed
E
Vincristine
F
Paclitaxel
G
Doxorubicin
H
Epirubicin
I
Bleomycin
J
Cisplatin
K
Oxaliplatin
L
ifosfamide
M
Mitoxantrone
N
Fludarabine
O
Erlotinib
P
AZD4547
Q
DMOG_165
R
Gefitinib_1010
S
Olaparib
T
Rucaparib
U
Talazoparib
V
Niraparib
W
Imatinib
X
Cetuximab

**Supplementary Figure 11. Comparisons of drug sensitivities and identification of the potential targeted compounds among the BC TIDE subtypes**. **(A)** Submap analysis reflects the sensitivity of the TIDE subtypes to the targeted treatments in the BC patients. **(B-X)** Comparisons of sensitivities of the TIDE subtypes to clinically recommended drugs: methotrexate **(B)**, gemcitabine **(C)**, pemetrexed **(D)**, vincristine **(E)**, paclitaxel **(F)**, doxorubicin **(G)**, epirubicin **(H)**, bleomycin **(I)**, cisplatin **(J)**, oxaliplatin **(K)**, ifosfamide **(L)**, mitoxantrone **(M)**, fludarabine **(N)**, erlotinib **(O)**, AZD4547 **(P)**, DMOG_165 **(Q)**, gefitinib **(R)**, olaparib **(S)**, rucaparib **(T)**, talazoparib **(U)**, niraparib **(V)**, imatinib **(W)** and cetuximab **(X)**. $^{*}p < 0.05$, $^{**}p < 0.01$, $^{***}p < 0.001$, $^{****}p < 0.0001$; ns, no significance.

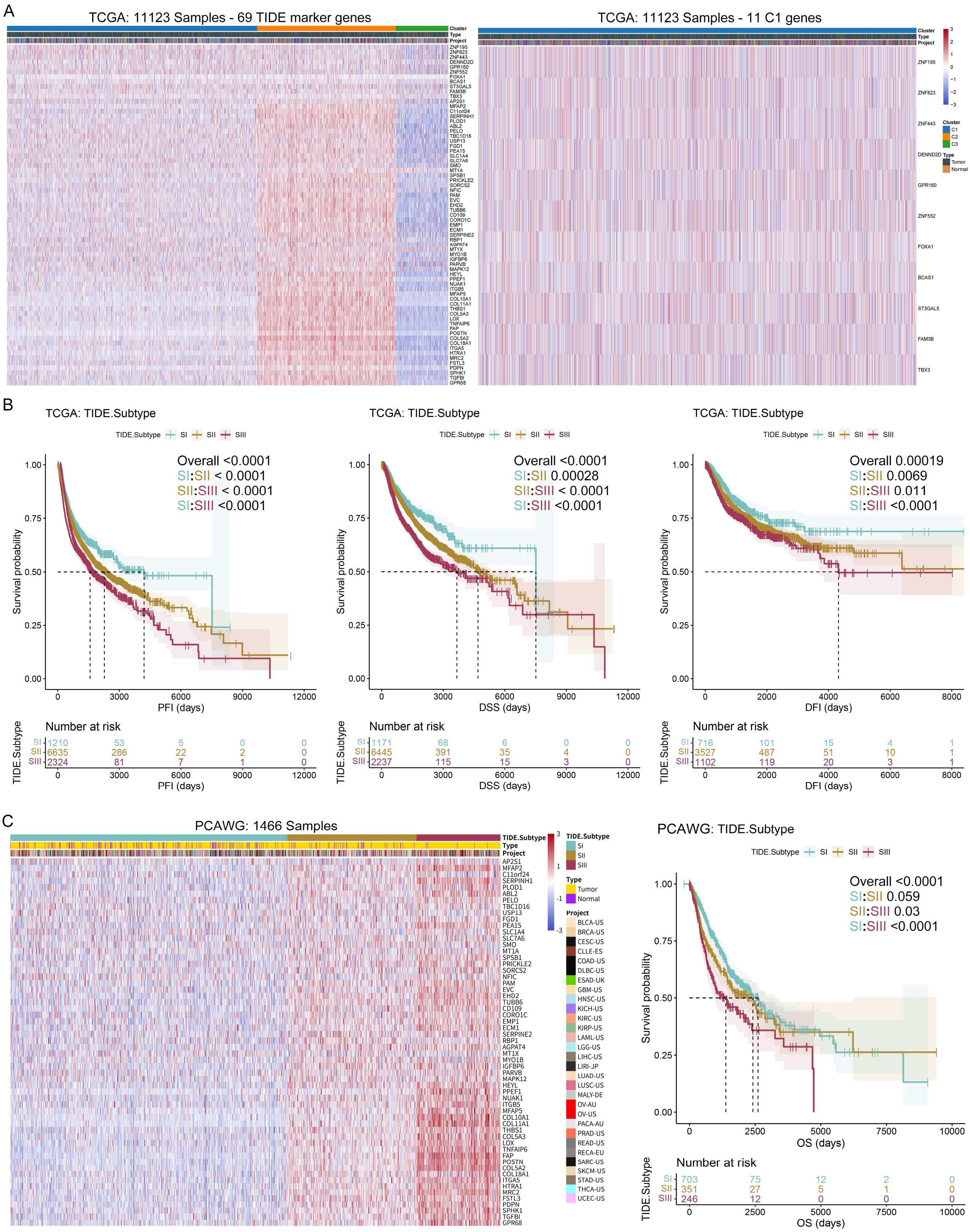
A
TCGA: 11123 Samples - 69 TIDE marker genes
TCGA: 11123 Samples - 11 C1 genes
Cluster
Type
Project
C1
C2
C3
Tumor
Normal
ZNF195
ZNF823
ZNF443
DENND2D
GPR160
ZNF552
FOXA1
BCAS1
ST3GAL5
FAM3B
TBX3
B
TCGA: TIDE.Subtype
TIDE.Subtype SI SII SIII
Overall <0.0001
SI:SII < 0.0001
SII:SIII < 0.0001
SI:SIII <0.0001
Survival probability
PFI (days)
Number at risk
SI 1210 53 5 0 0
SII 6635 286 22 2 0
SIII 2324 81 7 1 0
Overall <0.0001
SI:SII 0.00028
SII:SIII < 0.0001
SI:SIII <0.0001
DSS (days)
SI 1171 68 6 0 0
SII 6445 391 35 4 0
SIII 2237 115 15 3 0
Overall 0.00019
SI:SII 0.0069
SII:SIII 0.011
SI:SIII <0.0001
DFI (days)
SI 716 101 15 4 1
SII 3527 487 51 10 1
SIII 1102 119 20 3 1
C
PCAWG: 1466 Samples
TIDE.Subtype
Type
Project
PCAWG: TIDE.Subtype
Overall <0.0001
SI:SII 0.059
SII:SIII 0.03
SI:SIII <0.0001
OS (days)
SI 703 75 12 2 0
SII 351 27 5 1 0
SIII 246 12 0 0 0

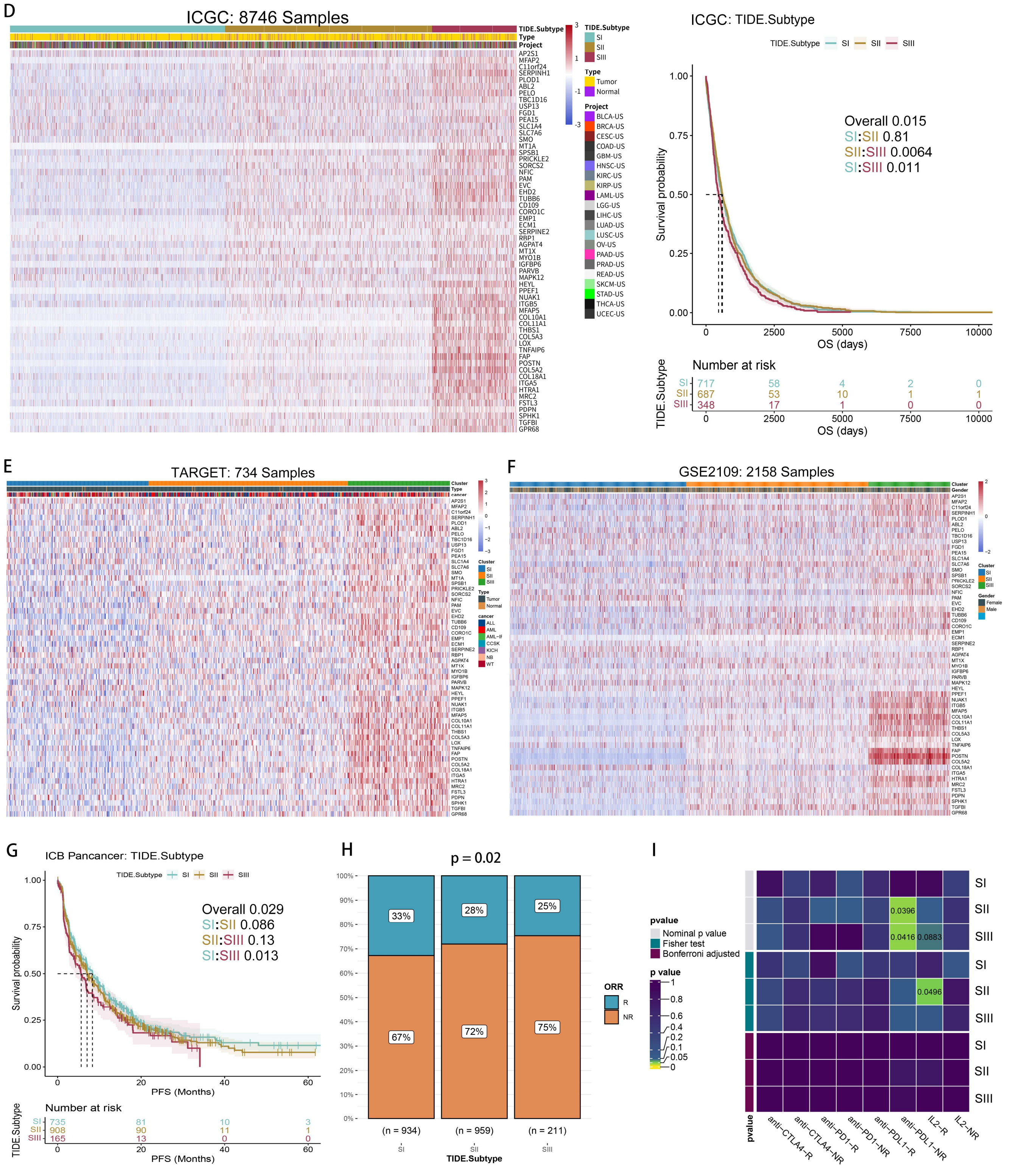


**Supplementary Figure 12. Conservations of the TIDE subtypes in pan-tumors** (related to Figure 8). **(A)** Unsupervised hierarchical clustering based on the 69 TIDE marker genes classified pan-tumor samples from TCGA into three subtypes.11 C1 genes were approximately equally expressed among the TIDE subtypes **(left)**. Unsupervised clustering analysis using the 11 C1 genes failed to cluster the pan-cancer samples **(right). (B)** K-M analysis showing significant differences in PFI, DSS and DFI among the TIDE subtypes of TCGA pan-tumors. **(C-F)** Unsupervised hierarchical clustering based on the 58 C2 genes classified pan-tumor samples from PCAWG **(C)**, ICGC **(D)**, TARGET **(E)** and GSE2109 **(F)** into three TIDE subtypes. K-M analysis shows distinct prognosis of the TIDE subtypes of PCAWG **(C-right)** and ICGC **(D-right)**. **(G)** K-M analysis shows distinct prognosis of the TIDE subtypes in the ICB-treated pan-tumor cohort. **(H)** Stacked histogram showing the differences of pan-tumor ICB responses among the TIDE subtypes. **(I)** Submap analysis showing the sensitivity of the pan-tumor TIDE subtypes to ICB therapy.

### Supplementary Methods

#### Study Design

The design of this study is as follows:

(1) Evaluation of tumor immune dysfunction and exclusion (TIDE) status and its relationship with clinicopathological and molecular features of bladder cancer (BC): We used the Tumor Immune Dysfunction and Exclusion (TIDE) algorithm[40] to evaluate the TIDE status and calculate the TIDE scores of samples in eight bulk RNA-seq datasets of BC[1-5]. Then, we explored the association between TIDE scores and clinicopathological features (such as pathological TNM staging, age, weight, race, gender, and follow-up information) and molecular features (such as tumor mutational burden [TMB], somatic copy number alterations) of BC.

(2) TIDE subtyping of BC: First, we identified 69 TIDE marker genes through correlation analysis and univariate COX analysis. Then, we divided BC patients into three subtypes using unsupervised consensus clustering[41] based on these TIDE marker genes.

(3) Characterization of clinicopathological and molecular features of TIDE subtypes: First, we used two additional algorithms - Unsupervised hierarchical clustering and Non-negative Matrix Factorization (NMF)[42] - based on 69 TIDE marker genes to classify BC samples, to test the stability of the subtypes. Next, we characterized the differences in clinical and pathological features, molecular features, tumor immune microenvironment (TIME), functional annotations and signaling pathways, as well as drug sensitivity of three TIDE subtypes.

(4) Analysis of the pan-tumor landscape of the TIDE subtypes: First, we performed unsupervised hierarchical clustering on five pan-tumor datasets based on 69 TIDE marker genes, dividing each dataset into three TIDE subtypes. Next, we validated the sensitivity differences of the three TIDE subtypes to immune checkpoint blockade (ICB) therapy using a bulk RNA-seq pan-tumor cohort of baseline samples treated with ICB.

### Datasets collection

Five bulk RNA-seq datasets[1-5] and one scRNA-seq dataset[39] of BC, five bulk-RNA-seq cohorts of pan-tumors, one bulk RNA-seq cohort of pan-tumors treated with ICB[6-38], and somatic mutation and CNA data from TCGA-BLCA were collected in this study. Bulk RNA-seq datasets of BC and follow-up information of TCGA-BLCA were collected from TCGA, Gene Expression Omnibus (GEO) and UCSC Xena. The CNA and somatic mutation data were downloaded from TCGA using the TCGAbiolinks package (v2.25.3). Single-cell RNA-seq dataset of BC was collected from the Mendeley Data (https://data.mendeley.com/datasets/7yb7s9769c/1). Pan-cancer cohorts were collected from UCSC Xena and GEO. The pretreatment bulk RNA-seq cohort of pan-tumors treated with ICB was obtained from multiple databases (**Table S1**). This cohort included 2641 samples from 36 datasets of 12 tumor types. The database of MSigDB[43] (v2023.1.Hs, http://www.gsea-msigdb.org/gsea/index.jsp) and STRING[44] (v11.5, https://cn.string-db.org/) were also collected in this study. Details and sources for all datasets are listed in **Table S1**.

#### Real-world blader tumor samples collection

After obtaining patient consent and approval from the institutional research ethics committee, we collected 31 surgical resection samples of bladder tumors from 20 patients who hospitalized in the Department of Urology, Shanghai Sixth People's Hospital. **Table S2** lists detailed information of the patients. For all resected tumor samples, we invited pathology experts to review the samples for accurate pathological diagnosis of the patients. All tumor samples were collected within 5 minutes after excision and immediately placed in liquid nitrogen for preservation. Subsequently, they were operated for bulk RNA sequencing (LY dataset) to validate the TIDE subtypes.

#### Bulk RNA sequencing and processing

RNA quality was measured with Bioanalyzer 2100 system and RNA Nano 6000 Assay Kit. mRNA was purified from total RNA with magnetic beads with poly-T oligos. mRNA was fragmented with divalent cations and high temperature. First and second strand cDNA were made with random primers, M-MuLV Reverse Transcriptase (RNase H-), DNA Polymerase I and RNase H. cDNA ends were polished and adenylated. Adaptor with hairpin loop structure was ligated to cDNA. cDNA fragments of about 370~420 bp were selected with AMPure XP system. PCR was done with Phusion High-Fidelity DNA polymerase, Universal PCR primers and Index (X) Primer. PCR products were cleaned with AMPure XP system and library quality was checked with Agilent Bioanalyzer 2100 system. Index-coded samples were clustered with cBot Cluster Generation System and TruSeq PE Cluster Kit v3-cBot-HS (Illumia). Libraries were sequenced on Illumina Novaseq platform and 150 bp paired-end reads were obtained. Raw data of fastq format were processed with fastp software. Reads with adapter, ploy-N and low quality were removed. Clean data were obtained and Q20, Q30 and GC content were calculated. Reference genome and gene model annotation files were downloaded from genome website. Reference genome index was built with Hisat2 v2.0.5. Clean reads were aligned to reference genome with Hisat2 v2.0.5. Gene model annotation file was used by Hisat2 to make splice junctions database and get better mapping result. Mapped reads were assembled in reference-based way with StringTie (v1.3.3b). StringTie used new network flow algorithm and optional de novo assembly step to make and quantify full-length transcripts for each gene locus. Reads numbers for each gene were counted with featureCounts v1.5.0-p3. FPKM of each gene was calculated based on gene length and reads count.

#### Single-cell RNA-seq dataset processing

Single-cell RNA-seq dataset was processed using uniform methods and standards with the Seurat package[45] (v4.2.1). Cells with less than 200 genes or more than 20% mitochondrial genes were removed, as well as features detected in less than 3 cells. Normalization was performed using Seurat's defaults and 2000 hypervariable features were selected for downstream analysis. The sample integration and batch effect removal were performed using the anchor-based canonical correlation analysis (CCA) method, followed by data normalization using the ScaleData function. The integrated RNA assay was used for dimensionality reduction and clustering, and principal component analysis (PCA) was performed using the RunPCA function. Louvain clustering was done with 30 PCs and resolution = 1, and the results were visualized in two-dimensional space using the uniform manifold approximation and projection (UMAP). To identify marker genes for each cluster, differential expression (DE) analysis was conducted using the FindAllMarkers function. SingleR package[46] (v2.0.0) was used to annotate all clusters with relevant marker genes reported in the literature. Single-cell pseudotime and trajectory analysis was implemented with the Monocle 2 package (v2.22.0)[47].

#### Differential expression (DE) analysis

Marker genes for each Louvain cluster were identified using the FindAllMarkers function[45]. For bulk RNA-seq datasets, DE analysis was performed using the scCODE package (v1.2.0.1)[48]. Genes with Bonferroni FDR-corrected p-values < 0.05 and Detected_Times ⩾ 3 were considered as DE genes.

#### Tumor immune dysfunction (TID) and tumor immune exclusion (TIE) analysis

The computational framework developed by Liu et al., called Tumor Immune Dysfunction Exclusion (TIDE), is used to evaluate the TID and TIE status of cancer patients[40]. This algorithm evaluates TIDE scores based on two cancer immune escape mechanisms: the promotion of T cell dysfunction in tumors exhibiting significant infiltration of cytotoxic T lymphocytes (CTL), and the inhibition of T cell infiltration in tumors characterized by a low level of CTL[40]. Higher TIDE scores indicate greater TID and TIE levels. We used the TIDE package (v1.3) in Python software to assign TIDE score for each sample.

In the K-M analysis, the optimal cutoff point was calculated using the built-in surv_cutpoint function in the Survminer package (v0.4.9) to divide the patients into high and low TIDE groups.

#### Protein–protein interaction network analysis (PPI)

PPI network of TIDE marker genes were constructed using the STRING database[44] (https://cn.string-db.org/).

#### Clustering analysis

Consensus clustering was carried out using the ConsensusClusterPlus package[41] (v1.58.0). To improve the reliability of our clustering, we repeated the process 1000 times, randomly selecting 80% of the samples each time. To ensure the accuracy of our classification results, we employed two additional techniques: unsupervised hierarchical clustering and non-negative matrix factorization (NMF). When performing unsupervised hierarchical clustering, we applied hclust to the normalized data using default parameters, which perform agglomerative clustering using the complete linkage method. We then visualized the results using a dendrogram, which represents the hierarchical structure of the clusters. To specify a desired number of clusters, we used the cutree function to cut the dendrogram. Meanwhile, we employed the NMF algorithm from the NMF package[42] (v0.24.0), which decomposes the matrix and performs 100 repetitions for stable, unsupervised clustering.

The optimal number of clusters was comprehensively determined by the consensus heatmap, the cumulative distribution function (CDF) curves, and the proportion of ambiguous clustering algorithm (PAC)[49] (**Figure S5A**).

### Signaling pathway analysis

Download the reference gene-sets for GSEA analysis, including GOBP, Hallmark, KEGG, and Reactome, from the Molecular Signatures Database[43] (MSigDB, http://www.gsea-msigdb.org/gsea/index.jsp). The Investigate Gene Sets tool (http://www.gsea-msigdb.org/gsea/msigdb/annotate.jsp) can be used to identify the overlapping gene-sets of the submitted gene-lists in MSigDB.

Use the "ssgsea" method of the GSVA package[50] (v1.44.5) to perform single-sample gene set enrichment analysis (ssGSEA) and assign corresponding signature activity scores to each sample.

Evaluate the enrichment scores of signaling pathways using gene set variation analysis (GSVA) with the "gsva" method of the GSVA package[50]. Use the Limma package[51] to perform differential analysis on the enrichment scores, which can identify pathways significantly enriched in different TIDE subtypes.

Perform gene set enrichment analysis (GSEA) on pre-ranked DE gene-lists using the clusterProfiler package[52] (v4.4.4), with reference to the GOBP, Hallmark, KEGG, and Reactome gene-sets.

Use the QIAGEN IPA software (IPA Winter Release December 2022, https://digitalinsights.qiagen.com/products-overview/discovery-insights-portfolio/analysis-and-visualization/qiagen-ipa/) to perform ingenuity pathway analysis (IPA) on identified DE genes of each TIDE subtype. The Pathways tool can expose the activation or inhibition status of canonical pathways in each subtype. The Disease and Function tool can reveal the correlation between the interested gene sets and biological functions or diseases.

### Somatic mutation and CNV analysis

We analyzed genomic characteristics and mutation spectrum of the TIDE subtypes by performing somatic mutation and CNV analyses. Somatic mutation data and CNV data of TCGA-BLCA were downloaded, followed by the identification of significant amplified or deleted genomic regions using GISTIC_2.0 (v6.15.30, https://cloud.genepattern.org/gp/pages/index.jsf). The G-score for each region was calculated for the amplitude and frequency of variations. Mutation types, frequencies, and CNVs were further analyzed and visualized using maftools package (v2.12.0)[53].

#### Survival Analysis

We performed survival analysis using the Survival package (v3.4.0) and the Survminer package (v0.4.9) based on the Kaplan-Meier method, and univariate and multivariate Cox regression analyses. The survival curve was plotted by the Kaplan-Meier method, and the log-rank test was applied to compare survival differences among the TIDE subgroups.

#### Evaluation of tumor immune microenvironment (TIME) patterns

We applied 54 immune-related signatures from Charoentong et al[54]. and Şenbabaoğlu et al[55]. to examine the TIME patterns of BC samples. The ssGSEA algorithm[50] was used to compute the active scores of these signatures. Consensus clustering[41] of the ssGSEA scores were used to classify BC samples into different immune subtypes. We also used the ESTIMATE package (v1.0.13)[56] to analyze the TME status using the transcriptome data. This package can estimate the levels of stromal cells, immune infiltration, and tumor purity. Moreover, we used the SCDC[57] and DECEPTICON (v1.0, https://github.com/Hao-Zou-lab/DECEPTICON) algorithms to evaluate the abundance of immunocytes in the tissue samples.

#### Evaluation of immunotherapy and targeted therapeutic efficacy

To evaluate the immunotherapy and targeted treatment responses among different TIDE subtypes, we employed unsupervised subclass mapping analysis[58] (SubMap, v4.0, https://cloud.genepattern.org/gp/pages/index.jsf). Submap is an unsupervised machine learning method to measure the similarity between TIDE subtypes and therapeutic efficacy, which helps us to determine the sensitivity to immunotherapy and targeted treatment in different groups[58]. Furthermore, we also validated this finding by analyzing a dataset of pan-tumors received ICB treatment.

#### Evaluation of drug sensitivity

The oncoPredict package[59] (v0.2) was used to evaluate drug sensitivity of TIDE subtypes using GDSC[60] (v8.4, https://www.cancerrxgene.org/) and CTRP[61] (v2, https://portals.broadinstitute.org/ctrp/) databases. The half-maximal inhibitory concentration (IC50) was used to measure drug sensitivity. The IC50 of drugs in BC patients was computed using ridge regression algorithm[59].

#### Identification of TIDE marker genes for TIDE subtyping.

We identified TIDE marker genes based on 8 bulk RNA-seq datasets of BC[1-6]. Spearman correlation analysis was performed between gene expression levels and TIDE scores of the datasets.

The selected genes were significantly correlated with TIDE scores (FDR < 0.05) called TIDE.Genes. Calculate the arithmetic mean of Spearman's R for each gene in the collected datasets. Genes were merged as TIDE.Sig if they were significantly correlated with TIDE score in at least four datasets and an average R not less than 0.3. The univariate COX analysis was performed on TIDE.Sig based on the survival data, and genes that were significantly associated with prognosis ($p < 0.05$) in at least four datasets were selected as TIDE marker genes.

#### Hematoxylin-eosin (H&E) staining and immunohistochemistry (IHC)

The collected BC tissue specimens were fixed in formalin and embedded in paraffin in the Pathology Department of the Shanghai Sixth People's Hospital. BC sections were cut from paraffin-embedded tissue blocks, soaked with 10% formaldehyde, and fixed on slides. The slides were incubated at 42°C overnight. For pathological evaluation, all sections were stained with H&E staining. For IHC analyses, sections were deparaffinized and rehydrated followed with antigen retrieval and serum blocking. They were incubated with primary antibodies at 4°C overnight, secondary antibodies at 37°C for 1 hour and Strept Avidin-Biotin Complex (SABC) at 37°C for 30 min. All sections were colored with DAB and countercolored with hematoxylin. The primary antibodies used were the Anti-KI67 mouse monoclonal antibody (1:200, C650056-0100, Sangon Biotech, Shanghai, China), Anti-CK5&6 mouse monoclonal antibody (1:200, C650026-0100, Sangon Biotech, Shanghai, China), Anti-CD44 mouse monoclonal antibody (1:200, D190741-0100, Sangon Biotech, Shanghai, China), and BeyoIHC™ Phosphohistone H3 (PHH3) Rabbit Monoclonal Antibody (1:200, AG8625, Byotime, Shanghai, China).

#### CT-based calculation of tumor volume

For mass-like tumors, we approximate them as half an ellipsoid, and we need to measure the long and short diameters of the largest cross-section of the tumors, count the number of layers where the tumor appears and record the CT slice thickness, the volume formula is:

$$\text{Volume} = \frac{\pi}{6} * L * S * H,$$

$$\text{H} = \text{number of layers} * \text{slice thickness},$$

where L is the long diameter, S is the short diameter, and H is the longitudinal diameter.

For the whole bladder infiltrating tumor, we approximate the bladder as an ellipsoid, and approximate the tumor volume as the bladder volume minus the bladder capacity, the formula is:

$$\text{Volume} = \frac{3\pi}{4} * L * S * H - \frac{3\pi}{4} * L' * S' * H',$$

$$\text{H or H}' = \text{number of layers} * \text{slice thickness},$$

Where L, S, H are the long, short, and longitudinal diameters of the bladder; L', S', H' are the same for the bladder cavity.

#### Statistical analysis

Parametric tests (independent sample t-tests and ANOVA) and non-parametric tests (Wilcoxon rank-sum test, Mann-Whitney U test, Kruskal-Wallis test, and Fisher's exact test) were used for normally and non-normally distributed or categorical variables, respectively. The Pearson correlation test and Spearman correlation test were utilized to evaluate the correlation between normally and non-normally distributed variables, respectively. Log-rank test was applied to compare survival differences between groups based on the Kaplan-Meier method. Statistical significance was considered at $P < 0.05$. Data visualizations and statistical analyses were conducted using R software (v4.2.1) and Python software (v3.10). Binomial 95% confidence intervals were used to report all confidence intervals (CIs).

11 Choueiri, T. K. et al. Immunomodulatory Activity of Nivolumab in Metastatic Renal Cell

Carcinoma. Clin Cancer Res 22, 5461-5471 (2016). https://doi.org:10.1158/1078-0432.Ccr-15-2839
12 Braun, D. A. et al. Interplay of somatic alterations and immune infiltration modulates response to PD-1 blockade in advanced clear cell renal cell carcinoma. Nat Med 26, 909-918 (2020). https://doi.org:10.1038/s41591-020-0839-y
13 Motzer, R. J. et al. Nivolumab for Metastatic Renal Cell Carcinoma: Results of a Randomized Phase II Trial. J Clin Oncol 33, 1430-1437 (2015). https://doi.org:10.1200/jco.2014.59.0703
14 Motzer, R. J. et al. Nivolumab plus Ipilimumab versus Sunitinib in Advanced Renal-Cell Carcinoma. N Engl J Med 378, 1277-1290 (2018). https://doi.org:10.1056/NEJMoa1712126
15 Mahoney, K. M. et al. Soluble PD-L1 as an early marker of progressive disease on nivolumab. J Immunother Cancer 10 (2022). https://doi.org:10.1136/jitc-2021-003527
16 Miao, D. et al. Genomic correlates of response to immune checkpoint therapies in clear cell renal cell carcinoma. Science 359, 801-806 (2018). https://doi.org:10.1126/science.aan5951
17 Ascierto, M. L. et al. The Intratumoral Balance between Metabolic and Immunologic Gene Expression Is Associated with Anti-PD-1 Response in Patients with Renal Cell Carcinoma. Cancer Immunol Res 4, 726-733 (2016). https://doi.org:10.1158/2326-6066.Cir-16-0072
18 Motzer, R. J. et al. Molecular Subsets in Renal Cancer Determine Outcome to Checkpoint and Angiogenesis Blockade. Cancer Cell 38, 803-817.e804 (2020). https://doi.org:10.1016/j.ccell.2020.10.011
19 Motzer, R. J. et al. Avelumab plus axitinib versus sunitinib in advanced renal cell carcinoma: biomarker analysis of the phase 3 JAVELIN Renal 101 trial. Nat Med 26, 1733-1741 (2020). https://doi.org:10.1038/s41591-020-1044-8
20 Liu, S. et al. Response and recurrence correlates in individuals treated with neoadjuvant anti-PD-1 therapy for resectable oral cavity squamous cell carcinoma. Cell Rep Med 2, 100411 (2021). https://doi.org:10.1016/j.xcrm.2021.100411
21 Phillips, D. et al. Immune cell topography predicts response to PD-1 blockade in cutaneous T cell lymphoma. Nat Commun 12, 6726 (2021). https://doi.org:10.1038/s41467-021-26974-6
22 van den Ende, T. et al. Neoadjuvant Chemoradiotherapy Combined with Atezolizumab for Resectable Esophageal Adenocarcinoma: A Single-arm Phase II Feasibility Trial (PERFECT). Clin Cancer Res 27, 3351-3359 (2021). https://doi.org:10.1158/1078-0432.Ccr-20-4443
23 Zhao, J. et al. Immune and genomic correlates of response to anti-PD-1 immunotherapy in glioblastoma. Nat Med 25, 462-469 (2019). https://doi.org:10.1038/s41591-019-0349-y
24 Kim, S. T. et al. Comprehensive molecular characterization of clinical responses to PD-1 inhibition in metastatic gastric cancer. Nat Med 24, 1449-1458 (2018). https://doi.org:10.1038/s41591-018-0101-z
25 Obradovic, A. et al. Immunostimulatory Cancer-Associated Fibroblast Subpopulations Can Predict Immunotherapy Response in Head and Neck Cancer. Clin Cancer Res 28, 2094-2109 (2022). https://doi.org:10.1158/1078-0432.Ccr-21-3570
26 Cho, J. W. et al. Genome-wide identification of differentially methylated promoters and enhancers associated with response to anti-PD-1 therapy in non-small cell lung cancer. Exp Mol Med 52, 1550-1563 (2020). https://doi.org:10.1038/s12276-020-00493-8
27 Jung, H. et al. DNA methylation loss promotes immune evasion of tumours with high mutation and copy number load. Nat Commun 10, 4278 (2019). https://doi.org:10.1038/s41467-019-12159-9
28 Patil, N. S. et al. Intratumoral plasma cells predict outcomes to PD-L1 blockade in non-small cell lung cancer. Cancer Cell 40, 289-300.e284 (2022). https://doi.org:10.1016/j.ccell.2022.02.002
29 Auslander, N. et al. Robust prediction of response to immune checkpoint blockade therapy in

metastatic melanoma. Nat Med 24, 1545-1549 (2018). https://doi.org:10.1038/s41591-018-0157-9
30 Pomeranz Krummel, D. A. et al. Impact of Sequencing Radiation Therapy and Immune Checkpoint Inhibitors in the Treatment of Melanoma Brain Metastases. Int J Radiat Oncol Biol Phys 108, 157-163 (2020). https://doi.org:10.1016/j.ijrobp.2020.01.043
31 Hugo, W. et al. Genomic and Transcriptomic Features of Response to Anti-PD-1 Therapy in Metastatic Melanoma. Cell 165, 35-44 (2016). https://doi.org:10.1016/j.cell.2016.02.065
32 Riaz, N. et al. Tumor and Microenvironment Evolution during Immunotherapy with Nivolumab. Cell 171, 934-949.e916 (2017). https://doi.org:10.1016/j.cell.2017.09.028
33 Liu, D. et al. Integrative molecular and clinical modeling of clinical outcomes to PD1 blockade in patients with metastatic melanoma. Nat Med 25, 1916-1927 (2019). https://doi.org:10.1038/s41591-019-0654-5
34 Gide, T. N. et al. Distinct Immune Cell Populations Define Response to Anti-PD-1 Monotherapy and Anti-PD-1/Anti-CTLA-4 Combined Therapy. Cancer Cell 35, 238-255.e236 (2019). https://doi.org:10.1016/j.ccell.2019.01.003
35 Chiappinelli, K. B. et al. Inhibiting DNA Methylation Causes an Interferon Response in Cancer via dsRNA Including Endogenous Retroviruses. Cell 169, 361 (2017). https://doi.org:10.1016/j.cell.2017.03.036
36 Lauss, M. et al. Mutational and putative neoantigen load predict clinical benefit of adoptive T cell therapy in melanoma. Nat Commun 8, 1738 (2017). https://doi.org:10.1038/s41467-017-01460-0
37 Garcia-Diaz, A. et al. Interferon Receptor Signaling Pathways Regulating PD-L1 and PD-L2 Expression. Cell Rep 19, 1189-1201 (2017). https://doi.org:10.1016/j.celrep.2017.04.031
38 Kraehenbuehl, L. et al. Pilot Trial of Arginine Deprivation Plus Nivolumab and Ipilimumab in Patients with Metastatic Uveal Melanoma. Cancers (Basel) 14 (2022). https://doi.org:10.3390/cancers14112638
39 Salomé, B. et al. NKG2A and HLA-E define an alternative immune checkpoint axis in bladder cancer. Cancer Cell 40, 1027-1043.e1029 (2022). https://doi.org:10.1016/j.ccell.2022.08.005
40 Jiang, P. et al. Signatures of T cell dysfunction and exclusion predict cancer immunotherapy response. Nat Med 24, 1550-1558 (2018). https://doi.org:10.1038/s41591-018-0136-1
41 Wilkerson, M. D. & Hayes, D. N. ConsensusClusterPlus: a class discovery tool with confidence assessments and item tracking. Bioinformatics 26, 1572-1573 (2010). https://doi.org:10.1093/bioinformatics/btq170
42 Brunet, J. P., Tamayo, P., Golub, T. R. & Mesirov, J. P. Metagenes and molecular pattern discovery using matrix factorization. Proc Natl Acad Sci U S A 101, 4164-4169 (2004). https://doi.org:10.1073/pnas.0308531101
43 Subramanian, A. et al. Gene set enrichment analysis: a knowledge-based approach for interpreting genome-wide expression profiles. Proc Natl Acad Sci U S A 102, 15545-15550 (2005). https://doi.org:10.1073/pnas.0506580102
44 Szklarczyk, D. et al. The STRING database in 2023: protein-protein association networks and functional enrichment analyses for any sequenced genome of interest. Nucleic Acids Res 51, D638-d646 (2023). https://doi.org:10.1093/nar/gkac1000
45 Hao, Y. et al. Integrated analysis of multimodal single-cell data. Cell 184, 3573-3587.e3529 (2021). https://doi.org:10.1016/j.cell.2021.04.048
46 Aran, D. et al. Reference-based analysis of lung single-cell sequencing reveals a transitional profibrotic macrophage. Nat Immunol 20, 163-172 (2019). https://doi.org:10.1038/s41590-018-0276-y

47 Qiu, X. et al. Reversed graph embedding resolves complex single-cell trajectories. Nat Methods 14, 979-982 (2017). https://doi.org:10.1038/nmeth.4402
48 Zou, J. et al. scCODE: an R package for data-specific differentially expressed gene detection on single-cell RNA-sequencing data. Brief Bioinform 23 (2022). https://doi.org:10.1093/bib/bbac180
49 Şenbabaoğlu, Y., Michailidis, G. & Li, J. Z. Critical limitations of consensus clustering in class discovery. Sci Rep 4, 6207 (2014). https://doi.org:10.1038/srep06207
50 Hänzelmann, S., Castelo, R. & Guinney, J. GSVA: gene set variation analysis for microarray and RNA-seq data. BMC Bioinformatics 14, 7 (2013). https://doi.org:10.1186/1471-2105-14-7
51 Ritchie, M. E. et al. limma powers differential expression analyses for RNA-sequencing and microarray studies. Nucleic Acids Res 43, e47 (2015). https://doi.org:10.1093/nar/gkv007
52 Yu, G., Wang, L. G., Han, Y. & He, Q. Y. clusterProfiler: an R package for comparing biological themes among gene clusters. Omics 16, 284-287 (2012). https://doi.org:10.1089/omi.2011.0118
53 Mayakonda, A., Lin, D. C., Assenov, Y., Plass, C. & Koeffler, H. P. Maftools: efficient and comprehensive analysis of somatic variants in cancer. Genome Res 28, 1747-1756 (2018). https://doi.org:10.1101/gr.239244.118
54 Charoentong, P. et al. Pan-cancer Immunogenomic Analyses Reveal Genotype-Immunophenotype Relationships and Predictors of Response to Checkpoint Blockade. Cell Rep 18, 248-262 (2017). https://doi.org:10.1016/j.celrep.2016.12.019
55 Şenbabaoğlu, Y. et al. Tumor immune microenvironment characterization in clear cell renal cell carcinoma identifies prognostic and immunotherapeutically relevant messenger RNA signatures. Genome Biol 17, 231 (2016). https://doi.org:10.1186/s13059-016-1092-z
56 Yoshihara, K. et al. Inferring tumour purity and stromal and immune cell admixture from expression data. Nat Commun 4, 2612 (2013). https://doi.org:10.1038/ncomms3612
57 Dong, M. et al. SCDC: bulk gene expression deconvolution by multiple single-cell RNA sequencing references. Brief Bioinform 22, 416-427 (2021). https://doi.org:10.1093/bib/bbz166
58 Hoshida, Y., Brunet, J. P., Tamayo, P., Golub, T. R. & Mesirov, J. P. Subclass mapping: identifying common subtypes in independent disease data sets. PLoS One 2, e1195 (2007). https://doi.org:10.1371/journal.pone.0001195
59 Maeser, D., Gruener, R. F. & Huang, R. S. oncoPredict: an R package for predicting in vivo or cancer patient drug response and biomarkers from cell line screening data. Brief Bioinform 22 (2021). https://doi.org:10.1093/bib/bbab260
60 Yang, W. et al. Genomics of Drug Sensitivity in Cancer (GDSC): a resource for therapeutic biomarker discovery in cancer cells. Nucleic Acids Res 41, D955-961 (2013). https://doi.org:10.1093/nar/gks1111
61 Rees, M. G. et al. Correlating chemical sensitivity and basal gene expression reveals mechanism of action. Nat Chem Biol 12, 109-116 (2016). https://doi.org:10.1038/nchembio.1986
